# Designed nutrient-gradient phenotyping enables predictive ionomics of tropical leafy greens

**DOI:** 10.64898/2026.09.16.751971

**Authors:** Itamar Shenhar, Miguel R. Pebes-Trujillo, Aravind Harikumar, Zhitong Zhao, Li Yi Tan, Magdiel Inggrid Setyawati, Jie He, Ittai Herrmann, Kee Woei Ng, Matan Gavish, Menachem Moshelion

**Affiliations:** School of Materials Science and Engineering, Nanyang Technological University, Singapore; The Robert H Smith Faculty of Agriculture, Food and Environment, Hebrew University of Jerusalem, Israel; Singapore-HUJ Alliance for Research and Enterprise (SHARE), Singapore; National Institute of Education - Natural Sciences & Science Education, Nanyang Technological University, Singapore; Nanyang Environment and Water Research Institute (NEWRI), Singapore; School of Computer Science and Engineering, Hebrew University of Jerusalem, Jerusalem, Israel

**Keywords:** Plant ionomics, nutrient-gradient phenotyping, tabular machine learning, functional phenotyping, precision fertigation, controlled-environment agriculture

## Abstract

Accurate prediction of edible-leaf mineral composition is challenging because tissue ionomes integrate nutrient supply, plant identity, growth dilution, water-balance regulation and cumulative environmental exposure. We combined a designed nutrient-gradient experimental framework with machine learning to predict leaf ionome using 1,163 Chinese spinach (*Amaranthus dubius*) and Chinese broccoli (*Brassica oleracea* Alboglabra Group) plants grown across 16 designed nutrient-gradient experiments in a semi-controlled tropical greenhouse. Targeted perturbations of N, P, K, Ca, Mg, Fe and Mo were combined with cumulative transpiration, integrated microclimate, shoot dry weight and laboratory quantification of 12 harvested-leaf mineral targets. The resulting ionome was broad, non-Gaussian and species dependent, with coordinated off-target shifts also evident for minerals whose supplied concentrations were held constant. We benchmarked AutoGluon and TabPFN using five-fold leakage-free, distribution-balanced grouped cross-validation, in which all biological replicates from each dosing condition were withheld together. Across previously unseen nutrient conditions, pooled out-of-fold R² ranged from 0.72 to 0.92, with broadly comparable performance between architectures. Learning curves showed rapid early gains for several targets but persistent condition-level generalization gaps for total reduced nitrogen, K, Mn and Zn. Model-agnostic Shapley additive explanations identified target-specific combinations of nutrient inputs, plant type, substrate, cumulative environment and physiological traits, while cross-model attribution agreement varied among minerals. Finally, reduced-feature models were tested on an independent grower-operated cohort of 99 plants. Calibration was closest for NO₃⁻, Mg and B, whereas Mn and Na showed substantial external bias. These results establish leakage-aware predictive ionomics as a robust framework for post-harvest lab-free crop mineral estimation while defining the calibration and sensing requirements for commercial deployment.

## 1. Introduction

As an integrated physiological phenotype, the plant ionome reflects the complete mineral-nutrient and trace-element composition of plant tissue and is shaped by the interplay among genotype, development, nutrient supply and environment. Ionomics was originally developed to quantify the inorganic components of biological systems at scale, enabling mineral profiles to be interpreted as system-level outputs rather than as isolated elemental measurements (Salt et al., 2008). In crop production, this perspective is particularly relevant because mineral accumulation is governed by interacting uptake, transport, storage and dilution processes, while the harvested tissue ultimately determines nutritional and commercial quality.

Tropical leafy vegetables present a demanding case for predictive ionomics. In crops such as *Amaranthus dubius* (Chinese spinach) and *Brassica oleracea* Alboglabra Group (Chinese broccoli), the harvested leaf is both the marketable organ and a complex, highly dynamic metabolic source–sink system. Its mineral profile integrates nutrient supply, crop identity, root-zone conditions, growth rate and microclimate over the entire cultivation cycle. Leafy vegetables are also prone to substantial variation in nitrate accumulation, which is influenced by nitrogen (N) supply, light environment and plant physiological status (Bian et al., 2020). Under tropical greenhouse conditions, high temperatures, rapid growth and short production cycles further increase the likelihood of marked variation in mineral concentrations among harvests. Predicting edible-leaf mineral composition therefore requires models that integrate fertilizer inputs with physiological and environmental histories rather than relying on nutrient supply alone.

A central challenge is that the leaf ionome is not a set of independent fertilizer-response traits. Macronutrient- and micronutrient-uptake pathways interact through shared transport systems, nutrient signaling, charge-balance competition and coordinated homeostatic regulation (Fan et al., 2021; Hanikenne et al., 2021). Consequently, perturbing one supplied mineral can shift the accumulation of other elements, producing multivariate ionomic responses that are difficult to capture with single-nutrient regression models. This interdependence is particularly important in precision fertilization, where optimizing one target mineral may unintentionally alter the broader edible-leaf mineral profile.

Addressing this challenge requires experimental and computational frameworks that treat the harvested-leaf ionome as an integrated multivariate phenotype rather than as a collection of independently optimized mineral endpoints.

Physiological fluxes add another layer of complexity. Although active transport mechanisms regulate specific ions, the delivery and distribution of many nutrients are coupled to plant water movement.

Calcium provides a well-established example: its uptake and distribution are closely linked to xylem transport and water flow to transpiring tissue (Gilliham et al., 2011). More generally, cumulative transpiration captures an integrated history of the soil–plant–atmosphere continuum (SPAC), which is expected to influence nutrient movement toward and within the shoot (Jones, 2013). Mineral concentration is also shaped by biomass accumulation. Rapid dry-matter production can dilute tissue mineral concentrations when biomass increases faster than nutrient accumulation, a phenomenon known as the growth-dilution effect (Jarrell and Beverly, 1981). These considerations motivate the inclusion of cumulative transpiration and shoot dry weight as physiological anchors for predicting harvested-leaf mineral concentration.

Machine learning (ML) provides a means of modeling these nonlinear, multivariate relationships, but its application to plant mineral nutrition has largely focused on individual traits, such as biomass, nitrogen status or transpiration, rather than on the complete ionome. More broadly, ML is increasingly used in plant science and breeding to connect heterogeneous predictors with complex phenotypes across biochemical, physiological and agronomic scales (van Dijk et al., 2021). Intensive ionomic datasets nevertheless present a distinctive computational problem: they are typically tabular, heterogeneous, biologically structured and limited in sample size because every observation requires destructive harvest and laboratory mineral quantification. These properties make ionomic data well suited to modern tabular-learning frameworks, provided that validation is rigorous and external transfer is explicitly tested.

Here, we benchmarked two state-of-the-art tabular-modeling approaches for predictive ionomics. AutoGluon-Tabular is an automated machine-learning framework that combines multilayer ensembles and stacked models to achieve strong performance on structured tabular datasets (Erickson et al., 2020). TabPFN is a tabular foundation model designed for small-to-medium-sized tabular datasets that uses prior-data-fitted inference to make predictions with limited task-specific training (Hollmann et al., 2025). These approaches represent complementary strategies: one constructs a custom ensemble from the observed dataset, whereas the other applies a foundation-model prior to tabular prediction. Their comparison allowed us to test whether predictive accuracy and feature-attribution structure were stable across distinct modeling paradigms.

In this study, we developed a predictive ionomics framework for tropical leafy-green production using 16 nutrient-gradient cultivation cycles conducted in a semi-controlled commercial greenhouse. The dataset combined targeted mineral inputs, plant type, substrate, cumulative environmental exposure, cumulative transpiration, shoot dry weight and laboratory measurements of 12 harvested-leaf mineral outputs. The design was intended to generate broad ionomic variation rather than a narrow steady-state nutritional range, enabling the models to learn both direct fertilizer effects and off-target mineral shifts. We hypothesized that perturbing a single supplied mineral would induce coordinated multi-ionomic changes across the harvested-leaf profile and that these structured responses could be learned by state of the art tabular models. Because internal cross-validation alone is insufficient to demonstrate generalizability to a different production domain, we also tested model transfer in an independent grower-operated commercial cohort that was completely withheld from training. This dual validation framework enabled us to assess both controlled-domain predictive accuracy and the limits of farmer-deployable ionomic monitoring. The study links designed mineral perturbations, continuous plant–environment phenotyping and modern tabular learning in a predictive framework for crop ionomics, with implications for precision fertilization, nutritional-quality monitoring and data-driven management of high-value horticultural systems.

## 2. Materials and methods

### 2.1 Experimental setup

The study was conducted in a semi-controlled commercial greenhouse at a vegetable farm in Singapore (Oasis Living Lab, Netatech Ltd, Singapore; 1°24′48.7″N, 103°43′10.4″E). The facility was covered with polyethylene film and equipped with rooftop and net-covered lateral openings. Four high-capacity fans provided ventilation under predominantly natural daylight conditions. Sixteen cultivation experiments were performed between 13 September 2023 and 2 January 2026. Each independent nutrient-gradient experiment represented one cultivation cycle from transplanting to harvest and was defined by plant type, substrate and target mineral (Table 1). Two commercial leafy-green crop types supplied by Netatech Ltd were studied: *Amaranthus dubius* (Chinese spinach; Fig. 1C) and *Brassica oleracea* Alboglabra Group (Chinese broccoli; Fig. 1D). Seeds were sown in peat-filled seedling trays and cultivated for 3–4 weeks before transplantation into 4-L pots (Model 18, Tefen Ltd, Nahsholim, Israel). The growing media were coarse silica sand (F2, Rock and Sand Industries Ltd, Singapore) or coco peat (250, Riococo Ltd, Sri Lanka). After transplantation, plants were grown for 4–5 weeks until physiological harvest maturity. For external commercial validation, parallel side-table units (9.3 m × 1.4 m × 0.15 m) were established adjacent to the experimental array to mimic nearby operational plots. The units contained commercial-grade coco peat and were managed by professional growers using local agronomic practices and a standard fertilizer formulation verified in triplicate (245.2 NO₃⁻, 2.97 NH₄⁺, 40.92 P, 258.6 K, 128.18 Ca, 45.73 Mg, 5.14 Fe, 0.60 Zn, 0.81 Mn, 0.78 B, 0.008 Mo, 42.28 Na and 95.66 S; mg L⁻¹). Plants of the same age and crop type were transplanted at 10-cm spacing and harvested concurrently with the experimental treatments, providing an independent assessment under operational production conditions (Fig. S1).

**Fig. 1.**
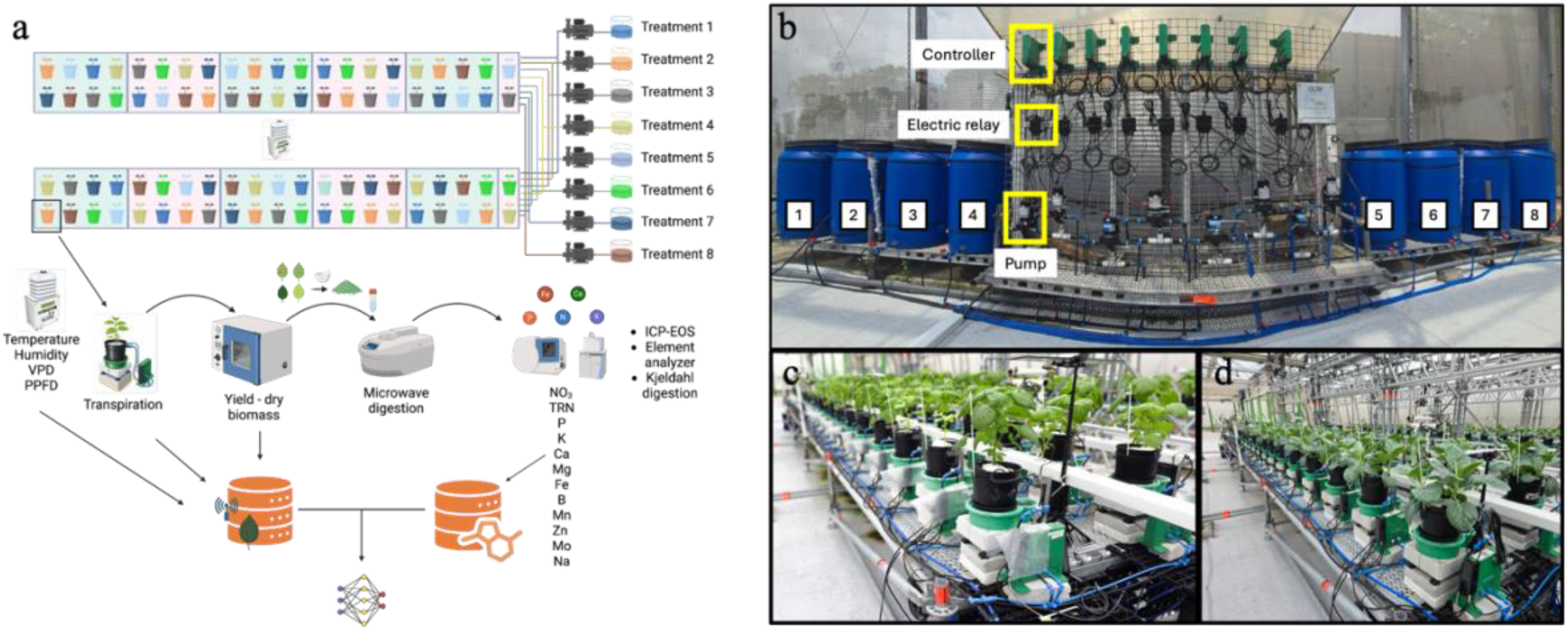
Experimental design, functional phenotyping platform and analytical workflow. (A) Randomized complete block design for eight nutrient-gradient treatments, continuous microclimate and physiological sensing, destructive postharvest biomass partitioning, tissue digestion, elemental profiling and integration into the machine-learning pipeline. (B) Automated multichannel fertigation system (Multi-Ferti, Plant-Ditech, Yavne, Israel), including the central controller, electrical relays and independent pumps for treatment reservoirs 1–8. (C) Chinese spinach at 21 days after transplanting (DAT) and (D) Chinese broccoli at 37 DAT on the high-throughput functional phenotyping platform. Each plant was coupled to an independent gravimetric lysimeter and microprocessor node that logged physiological and environmental data and controlled two irrigation valves through closed-loop feedback.

**Table 1.** Chronological timeline and experimental matrix for the 16 independent cultivation experiments used to construct the machine-learning dataset. The experiments spanned September 2023 to January 2026 in a semi-controlled tropical greenhouse and combined Chinese spinach and Chinese broccoli with coarse silica sand or coco peat. Each Experiment_ID denotes one nutrient-gradient cultivation cycle targeting N, P, K, Ca, Mg, Fe or Mo for downstream model training and validation.

| Experiment ID | Plant type | Substrate | Target mineral delivery | Cultivation dates |
| --- | --- | --- | --- | --- |
| 1 | *Amaranthus dubius* | Coarse silica sand | N | 13.09.2023–17.10.2023 |
| 2 | *Brassica oleracea* | Coco peat | N | 23.10.2023–29.11.2023 |
| 3 | *Amaranthus dubius* | Coarse silica sand | K | 13.12.2023–16.01.2024 |
| 4 | *Brassica oleracea* | Coco peat | K | 23.01.2024–27.02.2024 |
| 5 | *Amaranthus dubius* | Coarse silica sand | Mg | 04.03.2024–02.04.2024 |
| 6 | *Brassica oleracea* | Coco peat | Mg | 13.05.2024–18.06.2024 |
| 7 | *Amaranthus dubius* | Coarse silica sand | Fe | 25.06.2024–25.07.2024 |
| 8 | *Brassica oleracea* | Coco peat | Fe | 31.07.2024–07.09.2024 |
| 9 | *Amaranthus dubius* | Coarse silica sand | P | 20.09.2024–23.10.2024 |
| 10 | *Brassica oleracea* | Coco peat | P | 30.10.2024–02.12.2024 |
| 11 | *Amaranthus dubius* | Coarse silica sand | Ca | 13.12.2024–07.01.2025 |
| 12 | *Brassica oleracea* | Coco peat | Ca | 17.01.2025–20.02.2025 |
| 13 | *Amaranthus dubius* | Coco peat | N | 27.02.2025–26.03.2025 |
| 14 | *Amaranthus dubius* | Coco peat | K | 02.04.2025–02.05.2025 |
| 15 | *Amaranthus dubius* | Coco peat | P | 05.08.2025–04.09.2025 |
| 16 | *Brassica oleracea* | Coco peat | Mo | 26.11.2025–02.01.2026 |

The experimental design paired both crop types across the examined target minerals, with one exception. During the final molybdenum-gradient study (Experiment 16; Table 1), the *Amaranthus dubius* cohort developed severe disease symptoms during the late vegetative stage. To prevent disease-related effects from confounding the nutritional-stress analysis, the affected Chinese-spinach plants were excluded before destructive harvest. Experiment 16 therefore contained only *Brassica oleracea* Alboglabra Group.

### 2.2 Fertigation management and functional phenotyping platform

Custom nutrient solutions were formulated for each independent experiment to isolate specific ionomic responses. Within each cultivation cycle, one target mineral was varied across eight concentration levels, while the remaining nutrients were maintained at baseline concentrations except for counter-ion adjustments needed to preserve solution stability (Table S1). In some formulations, Na and S were introduced as counter-ions and therefore partly co-varied with the target-mineral gradient. Experiments were conducted on a high-throughput functional phenotyping platform (FFP; PlantArray, Plant-Ditech, Yavne, Israel) comprising 84 gravimetric lysimeter units for continuous, noninvasive monitoring of whole-plant physiological dynamics. Each fertigation treatment included 8–11 biological replicates arranged in a randomized complete block design across the platform (Fig. 1A). Differential nutrient delivery was controlled by the automated Multi-Ferti injection system (Plant-Ditech) coupled with high-precision diaphragm pumps (Shurflo 5050-1311-H011, Pentair, Minnesota, USA), which provided independent volumetric delivery to individual pots (Fig. 1B).

To maintain control over treatment delivery, multiparameter sensors (HYDROS 21, METER Group, Washington, USA) continuously monitored fertilizer-tank water depth, temperature and electrical conductivity (EC), while pH was measured with ruggedized glass electrodes (PHEHT, Ponsel-Aqualabo, Champigny-sur-Marne, France; Fig. S2). The central PlantArray system recorded sensor telemetry at 3-min intervals. Nightly irrigation operated under an automated closed-loop algorithm from 20:30 to 02:30. The first irrigation volume for each pot was adjusted according to that plant’s transpiration on the preceding day, followed by three equal 60-mL irrigation events to achieve the predetermined leaching fraction and return the substrate to field capacity each morning. A final 120-mL flushing event limited excessive nutrient accumulation in the root zone (Fig. S3).

### 2.3 Physiological and environmental monitoring

#### 2.3.1 Whole-plant water-use dynamics and cumulative transpiration

Continuous, noninvasive monitoring of soil–plant–atmosphere continuum (SPAC) hydraulics was used to characterize whole-plant water relations and functional physiological traits (Fig. S3). Diurnal transpiration and cumulative transpiration (CT) were calculated dynamically for each plant as described previously (Halperin et al., 2017; Dalal et al., 2020).

#### 2.3.2 Greenhouse microclimate sensing and cumulative environmental feature engineering

To capture spatial and temporal microclimatic heterogeneity within the semi-controlled greenhouse, a central meteorological station (WatchDog 2745, Spectrum Technologies Ltd, Bridgend, Wales) continuously recorded photosynthetic photon flux density (PPFD), air temperature, relative humidity (RH) and barometric pressure. Vapor pressure deficit (VPD) was calculated from these measurements. For model feature engineering, high-resolution observations from the diurnal photoperiod (07:00–19:00) were integrated over each cultivation cycle to generate four cumulative environmental variables: PPFD_SUM, VPD_SUM, Temp_SUM and RH_SUM (Table 2).

**Table 2.**
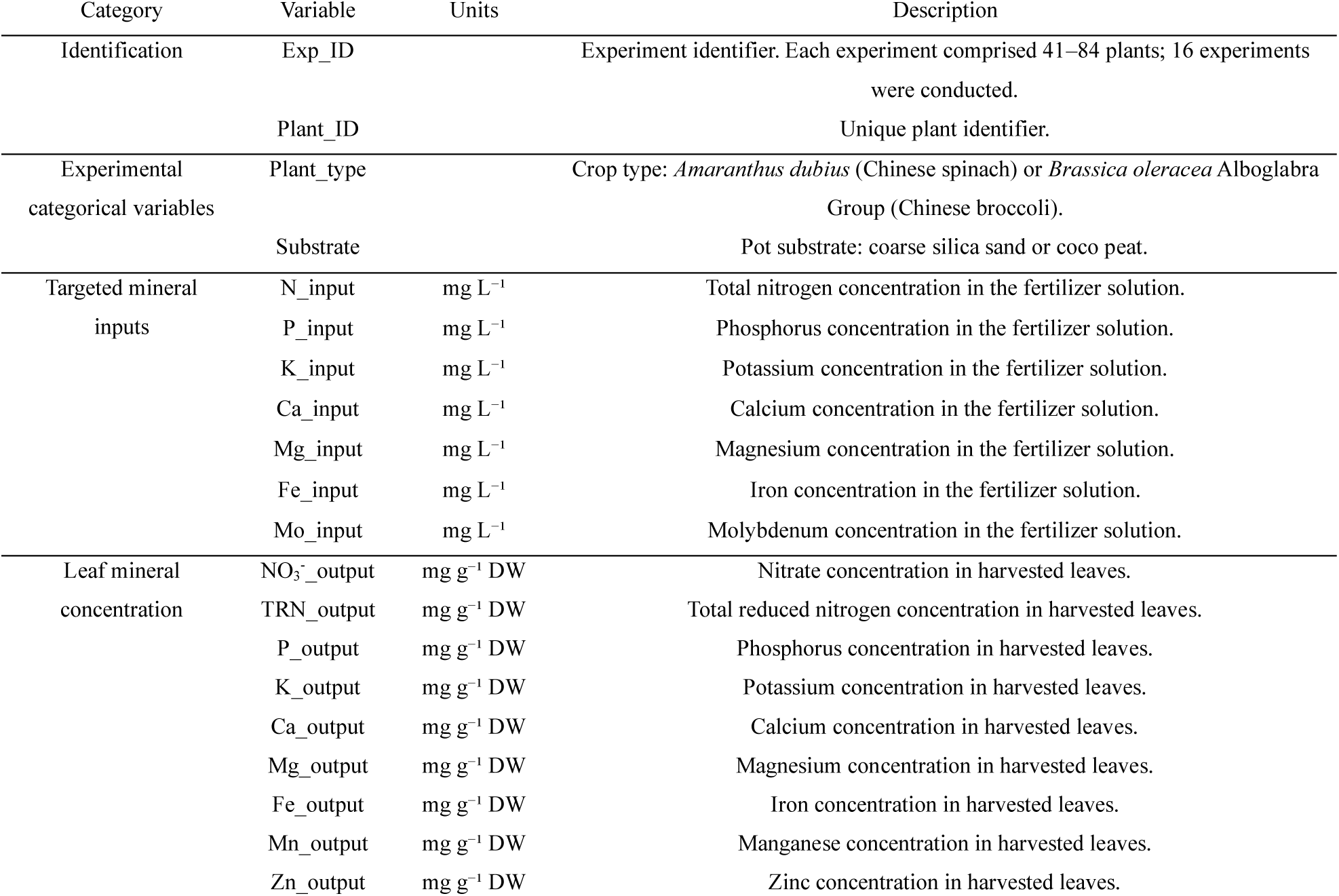
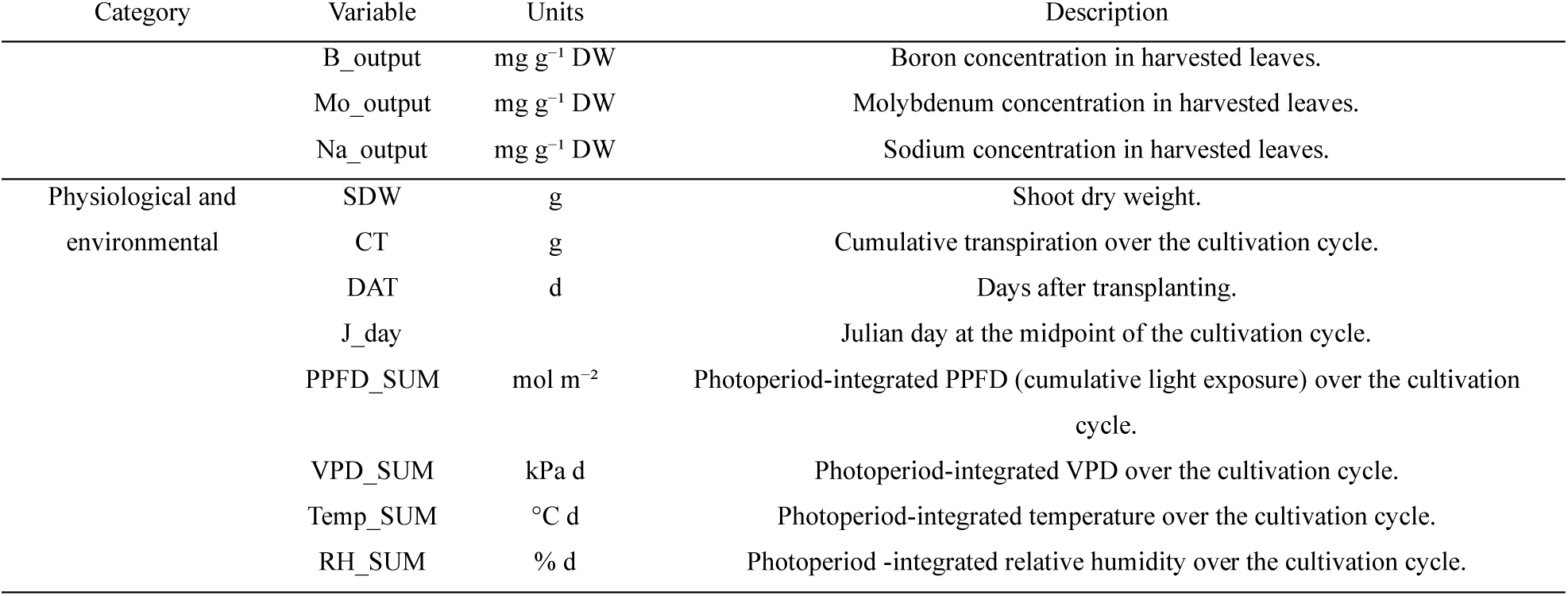
Multimodal dataset architecture and feature-space partitioning for predictive ionomic modeling. The 17 input variables describe crop identity and substrate, the seven target-mineral concentrations supplied through fertigation, and integrated physiological, environmental and temporal conditions. The 12 prediction targets are laboratory-measured harvested-leaf mineral concentrations.

#### 2.3.3 Deep-learning reconstruction of missing microclimate telemetry

A localized hardware failure during the potassium-gradient experiment with Chinese broccoli (Experiment 4) caused the loss of high-resolution internal microclimate telemetry. A deep-learning spatiotemporal reconstruction pipeline was implemented to retain this cultivation cycle without leaving its environmental features missing.

Parallel external microclimate time series, including air temperature, relative humidity (RH), rainfall (mm) and wind speed (knots), were obtained from a public meteorological station within 5 km of the experimental array (National Environment Agency, 2024). Estimated and theoretical solar radiation were retrieved through the Open-Meteo historical weather application programming interface (Zippenfenig, 2023). To model the nonlinear relationships between the external atmosphere and the semi-controlled greenhouse interior, a separate extended long short-term memory (xLSTM) recurrent neural network was constructed for each target variable. The networks were trained on data from the remaining successful cultivation cycles (349,525 observations), providing paired external-to-internal measurements for model development.

Target-specific xLSTM architectures and hyperparameters were optimized and validated as described in Table S2. The trained networks reconstructed the missing 3-min internal microclimate series for Experiment 4, and these estimates were integrated to obtain Temp_SUM, RH_SUM, PPFD_SUM and VPD_SUM (Fig. S4). Temperature, RH and VPD were reconstructed more accurately than PPFD, likely because localized tropical cloud cover produces faster and more spatially heterogeneous changes in radiation than in thermodynamic variables (Supplementary Note 1). The sensitivity of the ionomic results to this reconstructed experiment was evaluated separately (Fig. S5).

### 2.4 Postharvest processing and tissue ionomic profiling

#### 2.4.1 Organ-specific partitioning and biomass quantification

Immediately after destructive sampling, whole shoots were separated into leaf and stem fractions, with petioles included in the stem fraction. Tissues were dried in a forced-air oven (DHG-9920A, Bluepard Instrument, Shanghai, China) at 60 °C to constant mass. Whole-shoot dry mass was recorded as shoot dry weight (SDW; g).

#### 2.4.2 Sample processing and elemental analysis

After determination of dry mass, dried leaf tissue was separated from residual petiole material and homogenized to a fine powder. A 200-mg aliquot was digested by closed-vessel microwave-assisted acid digestion in an MPS 320 system (PerkinElmer Ltd, Massachusetts, USA) with 4 mL of 70% nitric acid (HNO₃) and 3 mL of Milli-Q water in MR-85 auto-venting TFM fluoropolymer vessels. After cooling, digests were diluted to 25 mL with deionized water. Potassium (K), phosphorus (P), calcium (Ca), magnesium (Mg), sodium (Na), manganese (Mn), zinc (Zn), molybdenum (Mo), iron (Fe) and boron (B) were quantified using an Avio 200 inductively coupled plasma optical emission spectrometer (ICP-OES; PerkinElmer Ltd). The instrument was operated at a radio frequency (RF) forward power of 1500W. Argon gas flows were maintained at 8 L min^−1^ for the plasma gas, 0.2 L min^−1^ for the auxiliary gas, and 0.6 L min^−1^ for the nebulizer gas. Samples were introduced into the system at a constant flow rate of 1.5 mL min^−1^ using a peristaltic pump. Instrument calibration, background correction and spectral processing were performed in Syngistix software (PerkinElmer Ltd). The resulting concentrations populated the corresponding mineral-output targets in the predictive dataset.

#### 2.4.3 Nitrogen fractionation: nitrate (NO₃⁻) and total reduced nitrogen (TRN)

Nitrogen was partitioned into inorganic nitrate and total reduced nitrogen fractions to characterize leaf N status. Foliar nitrate (NO₃⁻) was extracted and measured according to a published method (He et al., 2020). A 10-mg aliquot of dried, pulverized leaf tissue was extracted in 10 mL of Milli-Q water and agitated on an orbital shaker at 37 °C and 200 rpm for 2 h. The slurry was filtered through a 0.45-µm membrane under vacuum, and the filtrate was brought to 50 mL with acidified Milli-Q water. Nitrate was quantified with a QuikChem 8000 flow-injection analyzer (Lachat Instruments Inc., Milwaukee, USA) by measuring the magenta reaction product at 520 nm.

Total reduced nitrogen (TRN) was quantified by the classical Kjeldahl wet-digestion method adapted from Allen, 1989. Homogenized dry leaf tissue (50 mg) was digested with a standard catalytic Kjeldahl tablet in 5 mL of concentrated sulfuric acid (H₂SO₄) for 60 min at 350 °C. After mineralization and cooling, TRN was quantified by steam distillation and potentiometric titration with a Kjeltec 8400 analyzer (Foss Tecator AB, Höganäs, Sweden).

### 2.5 Machine-learning analysis

#### 2.5.1 Data preprocessing, feature engineering and dataset construction

High-frequency environmental and physiological time series recorded at 3-min intervals were processed and aggregated to construct the tabular machine-learning dataset. For each of the 1,163 plants (Plant_ID) across 16 cultivation cycles (Exp_ID), measurements were integrated over the plant-specific interval from transplantation to harvest to generate cumulative scalar features. These included PPFD_SUM, VPD_SUM, Temp_SUM and RH_SUM, together with cumulative transpiration (CT), thereby representing the longitudinal microclimatic and physiological history of each biological replicate.

Environmental variables were integrated over the full cultivation window rather than represented by a single midday value because mineral concentration at harvest reflects cumulative exposure to light, temperature and atmospheric water demand. This approach follows established cumulative-exposure indices, including daily light integral (DLI; Faust and Logan, 2018) for PPFD and thermal time or growing-degree days for temperature (Bonhomme, 2000; McMaster and Wilhelm, 1997), while extending the same time-integrated logic to VPD and RH because VPD regulates stomatal conductance, transpiration and water transport over prolonged periods in protected crops.

The integrated variables were aligned with static experimental descriptors and laboratory outputs to form the multidimensional feature matrix (Table 2). The 17-feature model input space comprised three groups: (1) two categorical variables, plant type and substrate; (2) seven continuous fertigation-input variables, N_input, P_input, K_input, Ca_input, Mg_input, Fe_input and Mo_input; and (3) eight physiological, environmental and temporal variables, CT, SDW, PPFD_SUM, VPD_SUM, Temp_SUM, RH_SUM, J_day and DAT. The prediction targets were the final concentrations (mg g⁻¹ DW) of 12 harvested-leaf mineral fractions: NO₃⁻, TRN, P, K, Ca, Mg, Fe, Mn, Zn, B, Mo and Na.

#### 2.5.2 Cross-validation strategy and external commercial validation

To evaluate predictive performance, stability and generalizability while eliminating information leakage between training and validation sets, we combined distribution-balanced grouped cross-validation with an independent external validation cohort.

For internal benchmarking on the primary experimental dataset, we implemented five-fold Distribution-Optimally-Balanced Stratified Cross-Validation (DOB-SCV) (Moreno-Torres et al., 2012; Zeng and Martinez, 2000) in the scikit-learn framework (Kohavi, 1995; Pedregosa et al., 2011). The grouping unit was the complete experimental dosing condition (125 conditions), defined by experiment and exact target-mineral concentration level. All biological replicates grown under the same nutrient-dose condition were therefore assigned together to either training or validation.

Within each experiment, dosing conditions were additionally classified into low-, mid- and high-dose tiers according to the rank of the active-gradient concentration, using a symmetric rule that assigned the minimum dose to the low tier and the maximum dose to the high tier. A single deterministic distribution- balancing procedure then allocated complete dosing conditions among the five folds in round-robin order within each experiment × dose-tier stratum, with the starting fold cyclically shifted by experiment and tier. This produced one fixed partition in which every fold represented the full experimental design.

The resulting partition satisfied three prespecified properties (Fig. S6). First, the folds were group disjoint: 0% of validation plants shared a dosing condition with any training plant. Second, all 16 experiments were represented in both the training and validation subsets of every fold. Third, dose-tier representation did not deviate from uniformity (χ² test, p = 1.00). Each validation fold contained 24–26 dosing conditions, corresponding to approximately 80% of plants for training and 20% for validation.

The absence of information leakage was tested empirically with two permutation controls. Random permutation of response labels reduced pooled out-of-fold coefficient of determination (R²) to approximately zero (mean, −0.18 across the 12 targets), as expected when no exploitable signal remains. In a condition-level null test, each dosing condition was assigned a random constant response. DOB-SCV yielded R² = −0.48 because held-out conditions were unpredictable, whereas a condition-leaking partition yielded R² = 0.94, demonstrating the memorization pathway removed by grouped assignment.

As a robustness check, the complete benchmarking pipeline was also run with an alternative leakage-free StratifiedGroupKFold scheme (five folds, dosing condition as the group and dose-quintile stratification). Target-wise performance was highly concordant between the schemes (Pearson’s r = 0.93).

StratifiedGroupKFold was systematically but only marginally more conservative (mean ΔR² relative to DOB-SCV = −0.02; paired t-test, p = 0.017), consistent with its weaker guarantee of experiment representation within each fold.

The internal validation therefore tested prediction for previously unseen dosing conditions rather than for additional plants from conditions already represented in training. AutoGluon and TabPFN used identical fold assignments for all model comparisons, learning curves and Shapley additive explanations (SHAP) analyses, ensuring a like-for-like comparison between architectures.

#### 2.5.3 Automated machine learning

To establish a robust performance baseline for leaf-mineral prediction, we evaluated AutoGluon v1.5.0, an automated machine-learning framework for high-performance tabular modeling (Erickson et al., 2020). Separate regression models were fitted to each of the 12 mineral targets with TabularPredictor, root mean squared error (RMSE) as the optimization metric and the best_quality preset. This configuration used a multilayer stack ensemble with bagging and stacking (three stack levels) across gradient-boosted decision trees (CatBoost, LightGBM and XGBoost), randomized tree ensembles (random forests and extremely randomized trees) and neural networks (FastAI and PyTorch implementations). AutoGluon’s weighted-ensemble procedure combined out-of-fold predictions from the base learners to generate each final model.

Each full-training-set fit was allocated a wall-clock budget of 3,600 s, and each reduced-size fit used for the learning curves was allocated 900 s, yielding 600 independent AutoGluon fits (12 targets × five folds × 10 training fractions). The budgets were not binding: across 12 targets and five folds, full-size fits required a median of 323 s and a maximum of 1,818 s. The training-subsample draw used random seed 42; AutoGluon’s internal stochastic components, including bagging partitions and neural-network initialization, used framework defaults. Full configuration details and target-specific ensemble summaries are provided in Tables S3 and S4.

#### 2.5.4 Tabular foundation model

In addition to the ensemble-based automated machine-learning approach, we evaluated TabPFN v3 with tabpfn Python package v8.0.3 as a tabular foundation-model regressor for plant ionomics. TabPFN is a prior-data-fitted network that treats supervised tabular prediction as an in-context inference problem within a pretrained transformer architecture (Hollmann et al., 2025). We instantiated the v3 regression checkpoint (tabpfn-v3-regressor-v3_default) and applied it to the same feature matrix, targets and cross-validation folds used for AutoGluon. A fixed sampling seed ensured identical training subsamples across the two frameworks.

The model used a 32-member inference ensemble based on input permutations and feature and target transformations, automatic inference precision and random seed 42. The setting ignore_pretraining_limits=True was used as a precaution: with 17 features and at most 930 training observations per fold, the dataset remained within the architecture’s pretrained operating range, and no subsampling was triggered. Full implementation settings are provided in Table S5.

#### 2.5.5 Model evaluation and learning curves

Predictive performance was evaluated using R², RMSE and mean absolute error (MAE). Learning curves quantified data-saturation behavior and train–validation generalization gaps for every target. Within each cross-validation split, the held-out validation fold remained fixed while the corresponding training fold was progressively subsampled from 10% to 100% of the available observations. At each nonzero fraction, AutoGluon and TabPFN were trained independently and evaluated on the subsampled training set and the fixed held-out fold, producing paired training and validation R² values for every model, target, fraction and fold. Mean trajectories and standard-deviation ribbons were calculated across the five folds. The magnitude and persistence of the train–validation separation were used to assess model fit, and a paired Student’s t-test evaluated whether the gap differed from zero for each model and training fraction.

#### 2.5.6 Model interpretability and feature-importance analysis

To connect predictive performance with physiological interpretation, we applied SHAP analyses to the AutoGluon 1.5 and TabPFN v3 models (Lundberg and Lee, 2017). SHAP assigns a local contribution to each feature relative to a model-specific background expectation. Because the architectures do not share a common model-specific explainer, we used the model-agnostic Kernel SHAP estimator (KernelExplainer), which requires only the prediction function and could therefore be applied identically to both models. For each of the 12 targets, we explained the model fitted to the first DOB-SCV training fold using the same 250 instances for both architectures (random seed 42). The background distribution was summarized by k-means clustering into five centroids (k = 5), and each instance used 2,048 coalition samples. TabPFN v3 attributions used the same 32-member ensemble as prediction. A separate sensitivity analysis found that rankings were invariant to TabPFN ensemble size (Spearman’s ρ = 0.97–0.99; Pearson’s r ≥ 0.998; identical top five feature sets for 8- and 32-member explanations on representative targets). Global importance was calculated as the mean absolute SHAP value for each feature and target, and cross-model agreement was summarized by R² between the two feature-importance vectors. This comparison tested whether attribution patterns were stable across the two modeling paradigms.

Global feature effects were visualized as beeswarm plots. Each point represented one of the 250 explained instances, positioned by its SHAP value and colored from low to high according to the corresponding feature value. To limit the influence of extreme values, colors were normalized within each feature to its 5th–95th percentile range. Features were ordered independently within each model by decreasing mean absolute SHAP value, and the eight highest-ranked features were displayed. AutoGluon and TabPFN panels for each target shared a common SHAP-value range, allowing attribution magnitudes to be compared while retaining model-specific ordering. Attributions were computed on the research computing cluster described in Section 2.5.8, using NVIDIA L40S GPUs for TabPFN and AMD EPYC CPU nodes for AutoGluon, with shap 0.51.0 and scikit-learn 1.7.2.

Supplementary robustness analyses tested whether cross-model agreement and feature rankings depended on the SHAP configuration. Four representative targets were selected: TRN (nitrogen metabolism), Ca (transpiration- and transport-linked), Fe (micronutrient behavior) and Mo (low abundance). Relative to a common baseline of 150 explained instances, 2,048 coalitions, k = 5 and the first DOB-SCV fold, we varied the coalition count (1,024 versus 2,048), background size (k = 5, 10 or 20), explained-instance sample (independent draws A and B) and cross-validation fold. Stability was measured using Spearman’s rank correlation coefficient (ρ) between per-feature mean absolute SHAP vectors (Table S6). Mean ρ across the four targets was ≥ 0.99 for coalition count, 0.95–0.98 for background size and ≥ 0.96 for the explained-instance sample in both models. Fold-to-fold stability was lower (ρ = 0.87–0.94), as expected because each grouped fold withheld a different set of dosing conditions, but remained substantial and comparable between architectures. These results support the numerical robustness of the reported rankings while leaving their biological interpretation hypothesis generating.

#### 2.5.7 Adaptation to and validation in the commercial cohort under structural feature missingness

To evaluate transfer beyond the controlled experimental platform, we tested both pipelines in the independent grower-operated cohort. The commercial side tables lacked gravimetric balances and the destructive biomass workflow; cumulative transpiration (CT) and shoot dry weight (SDW), two of the 17 training features, were therefore unavailable for all commercial plants. The other 15 features were available: seven fertigation-mineral inputs, plant type, substrate, Julian day, days after transplanting and four cumulative microclimate variables (PPFD_SUM, VPD_SUM, Temp_SUM and RH_SUM).

We addressed this feature-space gap by retraining AutoGluon 1.5 and TabPFN v3 from scratch on the controlled dataset using only the 15 variables available in the commercial cohort, with CT and SDW excluded from both training and inference. This parsimonious strategy was chosen instead of imputing the two unavailable covariates because it represents a conservative deployment scenario based only on data that growers can realistically provide. It also isolates transfer under commercial fertigation, high planting density and root-zone competition without introducing error from a separate reconstruction model.

External performance was summarized for each mineral target by multiplicative bias, defined as the ratio of the mean predicted concentration to the mean observed concentration; a value of 1.0 indicates an unbiased central estimate. This metric was preferred to a regression-fit statistic because the 99 commercial plants shared one fertigation formulation and substrate. The seven mineral-input variables and substrate therefore carried no within-cohort variation, and between-plant predictions were driven primarily by plant type, Julian day, days after transplanting and the four cumulative microclimate variables. R² would penalize failure to explain variation about which most inputs contained little or no information, whereas multiplicative bias directly tests absolute calibration in the deployment domain.

#### 2.5.8 Computational implementation

Analyses were executed on the Hebrew University of Jerusalem Research Computing Services (HURCS) cluster under the Slurm workload manager. The different computational profiles of the two frameworks were assigned to different resources. AutoGluon ran on dual-socket AMD EPYC 7662 nodes (128 cores per node; 32 cores allocated per task), whereas TabPFN inference ran on NVIDIA L40S GPUs with eight CPU cores per task for data handling and ensemble parallelization. The benchmark was parallelized at the target × fold level and distributed as 60 CPU tasks and 12 GPU tasks per cross-validation scheme, with results merged after completion.

Across the two cross-validation schemes, external validation, calibration analysis and SHAP attribution, the study used approximately 13,900 CPU core-hours and 14.6 GPU-hours. The software environment was pinned to Python 3.11, AutoGluon 1.5.0, tabpfn 8.0.3, scikit-learn 1.7.2, shap 0.51.0 and PyTorch 2.6.0+cu124.

### 2.6 Statistical analysis

Statistical analyses were performed in Python with SciPy and scikit-learn. Model performance was summarized for each target with the metrics described in Section 2.5.5 and pooled across the DOB-SCV folds. Learning-curve trajectories are reported as fold means with standard-deviation ribbons. At each training fraction, paired Student’s t-tests compared training and validation R² values independently for each model and mineral target across the five folds. Significance was denoted by *p ≤ 0.05, **p ≤ 0.01 and ***p ≤ 0.001.

For SHAP-based interpretation, global feature importance was summarized as the mean absolute SHAP value per feature, model and target, as described in Section 2.5.6. Cross-model attribution agreement was quantified by R² between the AutoGluon and TabPFN feature-importance vectors. Robustness across alternative explanation settings was assessed with Spearman’s rank correlation coefficient. External commercial-validation statistics were treated as deployment-domain calibration diagnostics rather than as internal model-selection criteria.

## 3. Results

### 3.1 The experimental ionome spans broad, non-Gaussian and species-dependent mineral landscapes

Before evaluating predictive performance, we examined whether the experimental design generated sufficient ionomic diversity for learning mineral responses beyond narrow commercial steady-state conditions. The dataset comprised 16 independent cultivation cycles conducted over more than two years in a semi-controlled tropical greenhouse, combining two leafy-green species, contrasting substrates, targeted mineral gradients, destructive biomass measurements, continuous physiological monitoring and post-harvest mineral profiling (Fig. 1). Each plant was represented by its fertigation inputs, plant identity, substrate, cumulative environmental exposure, cumulative transpiration, shoot dry weight and harvested-leaf mineral concentrations. We therefore first visualized the empirical distribution of the harvested leaf ionome across fertilizer regimes and plant types using kernel density estimation (KDE; Fig. 2).

**Fig. 2.**
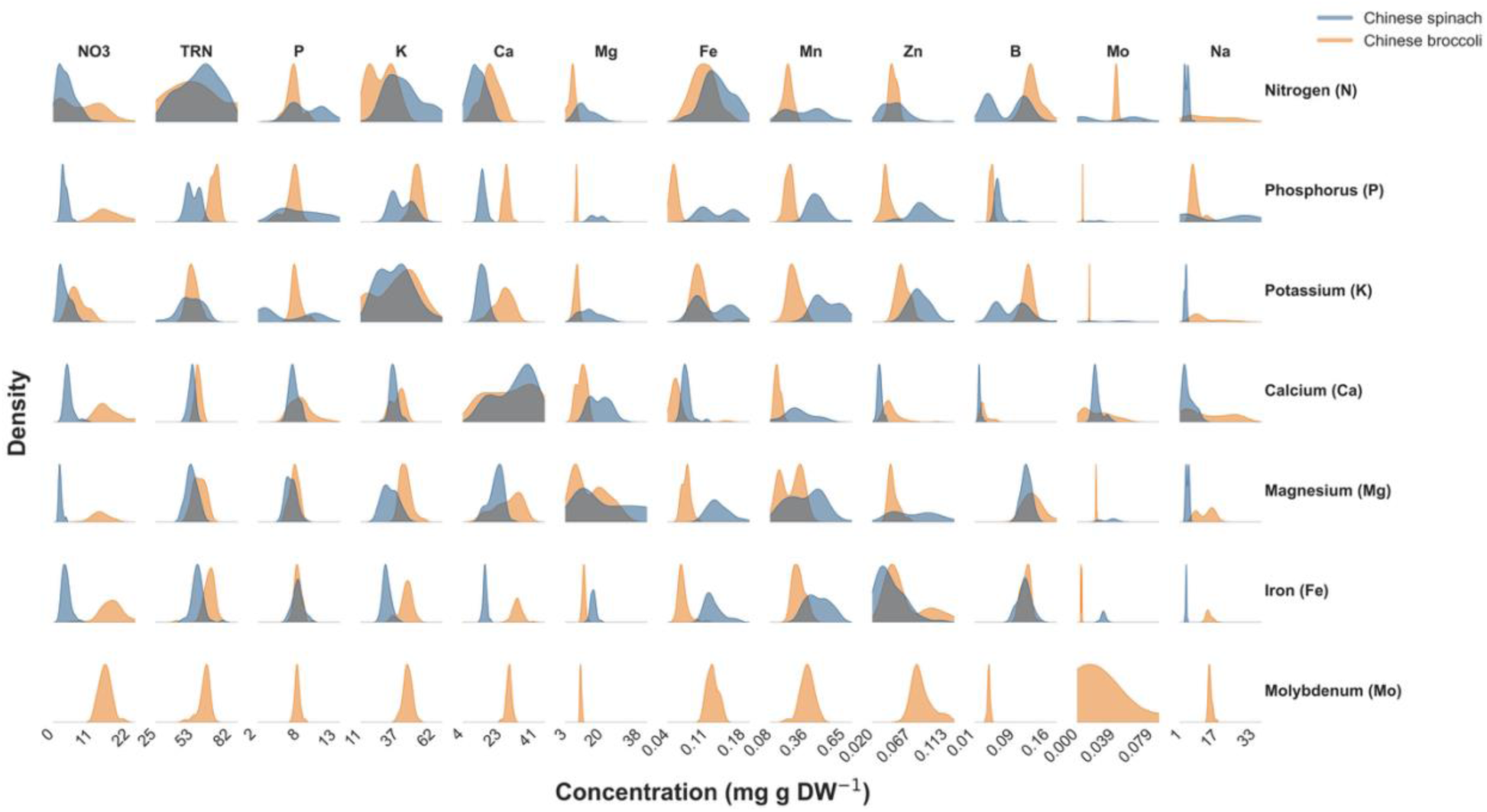
Kernel density estimates (KDEs) of harvested-leaf mineral concentrations across target-fertilizer experiments and plant types. Columns show the distributions of the 12 measured mineral targets (NO₃⁻, total reduced nitrogen, P, K, Ca, Mg, Fe, Mn, Zn, B, Mo and Na), and rows identify the mineral varied in each experiment. Curves are stratified by plant type (Chinese spinach and Chinese broccoli). The Mo experiment contains only Chinese broccoli because the diseased Chinese-spinach cohort was excluded before harvest, as described in Section 2.1.

The KDE matrix revealed broad, mineral-specific and frequently multimodal concentration distributions across the twelve predicted outputs: NO₃⁻, TRN, P, K, Ca, Mg, Fe, Mn, Zn, B, Mo, and Na. Some targets formed narrow density peaks, whereas others occupied broad or multimodal regions, indicating that the experimental matrix captured both tightly regulated mineral fractions and more plastic mineral pools.

Strong separation between Chinese spinach and Chinese broccoli was evident for several outputs, including the two nitrogen fractions, K, Ca, Mg and multiple micronutrients. Plant type therefore represented a major biological axis of ionomic variation rather than a minor categorical adjustment.

Targeted nutrient perturbations were also accompanied by shifts in non-target mineral distributions. These off-diagonal responses were most informative for elements whose supplied concentrations were held constant within each gradient experiment and are consistent with coordinated changes in uptake, transport, allocation, charge balance and growth dilution. Na was excluded from this crosstalk interpretation because its supplied concentration partly co-varied with several target gradients as a formulation counter-ion. Consequently, the observed Na patterns reflect a mixture of leaf-level response and engineered fertigation co-variation and cannot be interpreted solely as emergent ionomic coupling.

The two nitrogen outputs further illustrated the complexity of the prediction problem. NO₃⁻ and TRN displayed distinct distributional structures, consistent with their representation of inorganic nitrate storage and reduced-nitrogen pools, respectively. The Mo experiment formed an additional boundary case because it contained only Chinese broccoli after the diseased Chinese-spinach cohort was excluded before harvest. Overall, the dataset presented a structured, non-Gaussian and taxonomically heterogeneous prediction space in which harvested-leaf mineral concentration emerged from interacting nutrient, plant and environmental factors.

### 3.2 Leakage-free grouped cross-validation supports prediction across unseen dosing conditions

Predictive performance was evaluated using five-fold DOB-SCV, in which every biological replicate from a dosing condition was assigned to the same fold. Predictions from the five held-out validation folds were pooled so that every plant contributed exactly one out-of-fold prediction generated by a model that had not been trained on that plant’s dosing condition. All 16 experiments and low-, mid- and high-dose tiers remained represented in every fold. Under this leakage-free design, AutoGluon and TabPFN retained moderate-to-high performance across the complete ionomic panel (Fig. 3).

**Fig. 3.**
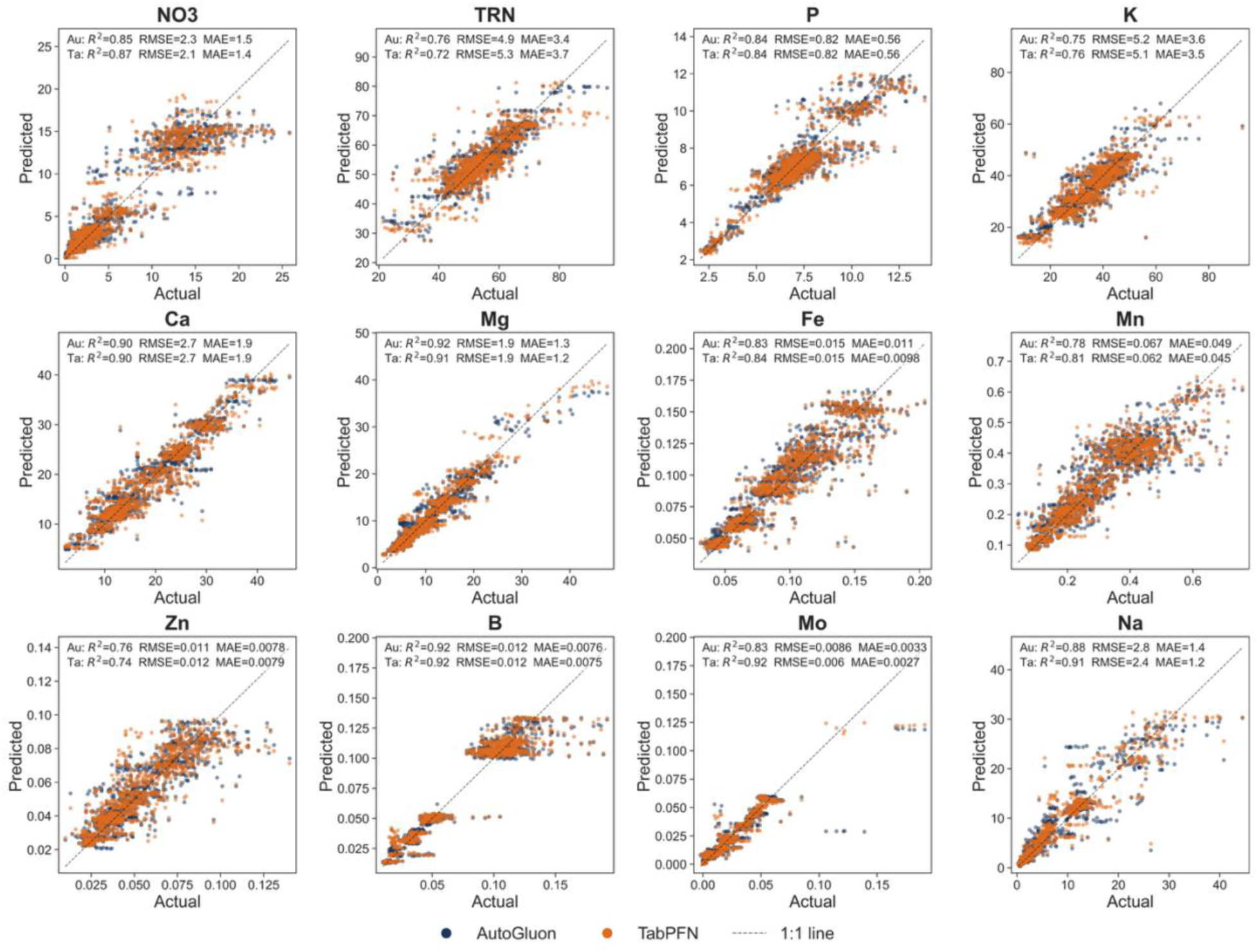
Leakage-free cross-validated prediction of 12 harvested-leaf mineral targets. Scatter plots compare observed and pooled out-of-fold predicted concentrations (mg g⁻¹ DW) from all five Distribution-Optimally-Balanced Stratified Cross-Validation (DOB-SCV) validation folds. Each plant contributes exactly one prediction, generated when its complete dosing-condition group was held-out from model training. Blue points show AutoGluon 1.5 predictions and orange points show TabPFN v3 predictions. For each model and mineral target, insets report coefficient of determination (R²), root mean squared error (RMSE) and mean absolute error (MAE) values were calculated from the same pooled observed-predicted pairs displayed in the corresponding panel; dashed lines denote 1:1 agreement.

Across the 24 model–target combinations, pooled out-of-fold R² values ranged from 0.72 to 0.92, with a mean of 0.84. B was predicted most consistently by both architectures (R² = 0.92), followed by Mg (R² = 0.92 for AutoGluon and 0.91 for TabPFN), Ca (R² = 0.90 for both) and Na (R² = 0.88 and 0.91, respectively). NO₃⁻ remained strongly predictable, with R² = 0.85 for AutoGluon and 0.87 for TabPFN. P and Fe reached R² values of approximately 0.84 in both architectures.

The largest architecture-specific difference occurred for Mo: TabPFN achieved R² = 0.92, RMSE = 0.0060 and MAE = 0.0027, whereas AutoGluon achieved R² = 0.83, RMSE = 0.0086 and MAE = 0.0033. Lower but still substantial performance was observed for Mn (R² = 0.78–0.81), K (R² = 0.75–0.76), Zn (R² = 0.74–0.76) and TRN (R² = 0.72–0.76). Residual dispersion was greatest at distributional extremes and in concentration ranges represented by relatively few held-out conditions.

Neither architecture consistently outperformed the other. TabPFN performed better for NO₃⁻, K, Fe, Mn, Mo and Na, whereas AutoGluon performed better for TRN, Mg and Zn; P, Ca and B were effectively tied. Their similar overall performance indicates that much of the predictive structure was shared, although the Mo result demonstrates an architecture-specific advantage for an individual target.

The permutation, condition-level null and alternative StratifiedGroupKFold controls described in Section 2.5.2 further supported the grouped partition, confirming that the reported performance did not arise from same-condition leakage or depend on one fold-construction algorithm.

### 3.3 Learning curves reveal rapid early gains and persistent target-specific generalization gaps

Learning curves were generated under the same grouped design by retaining each unseen-condition validation fold and progressively increasing the available training data. Both architectures extracted substantial predictive information from the first 10–30% of training observations, after which validation performance generally increased more gradually or approached an apparent plateau (Fig. 4). The relationship between training-set size and unseen-condition performance nevertheless differed markedly among targets.

**Fig. 4.**
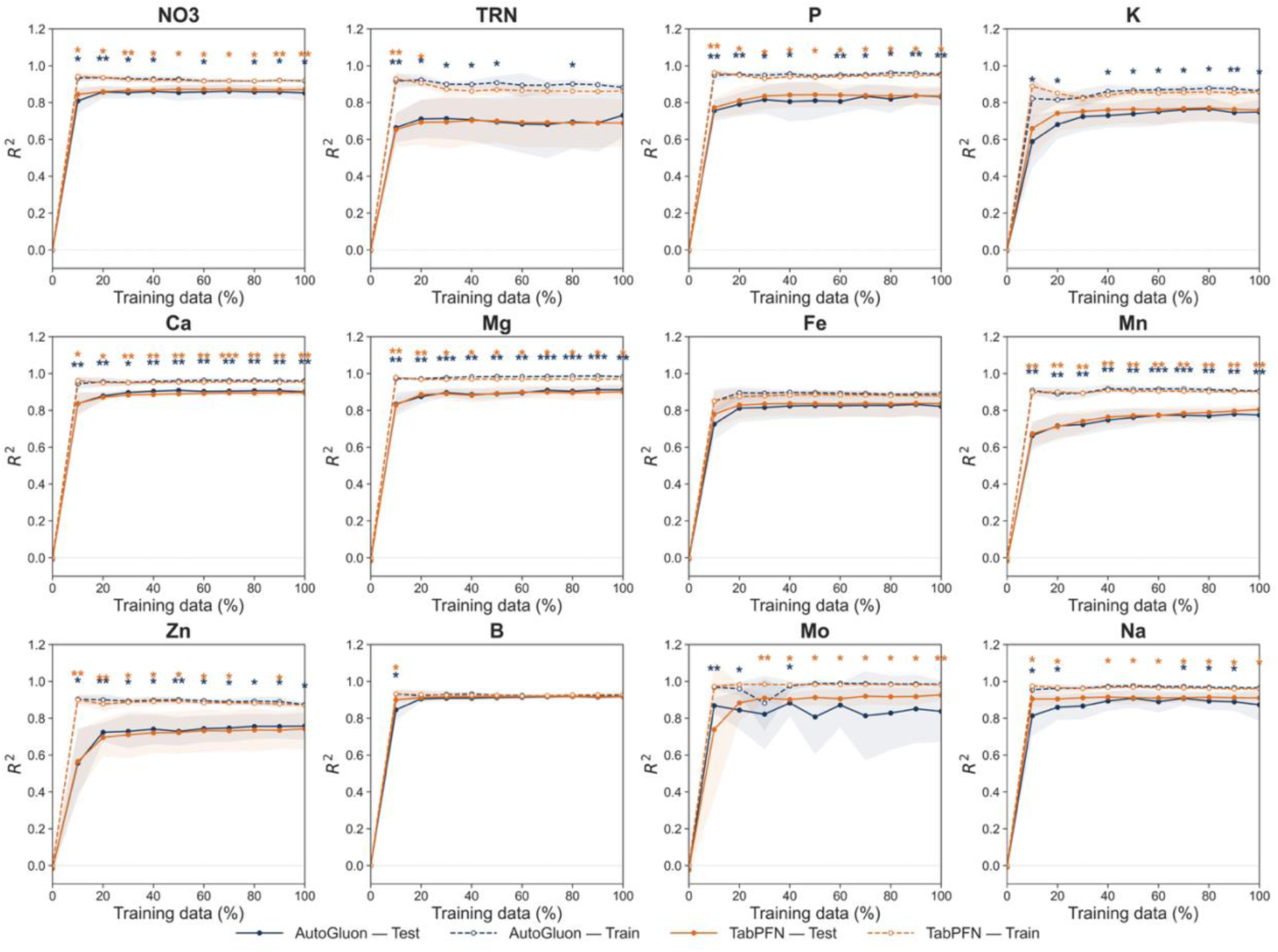
Learning curves for AutoGluon 1.5 and TabPFN v3 across the 12 harvested-leaf mineral targets. Coefficient of determination (R²) is shown as a function of the available training data (10–100%) under five-fold Distribution-Optimally-Balanced Stratified Cross-Validation (DOB-SCV). Dashed lines denote performance on the training subsets and solid lines denote performance on the fixed held-out folds. Points and shaded ribbons show fold means ± SD. Asterisks indicate paired Student’s t-tests of the train–validation difference across five folds: *p ≤ 0.05, **p ≤ 0.01 and ***p ≤ 0.001.

B showed the clearest convergence, with training and validation trajectories reaching similar performance at relatively low training fractions. NO₃⁻, P, Ca, Mg and Na also attained most of their final validation performance early, although modest train–validation separation persisted at larger fractions. Fe showed comparatively stable validation performance and a limited terminal gap.

TRN, K, Mn and Zn exhibited larger and more persistent train–validation separation. Their validation trajectories improved with additional data but remained below the corresponding training curves, indicating greater sensitivity to the diversity and coverage of the training conditions. Mo showed a model-specific pattern: TabPFN validation performance continued to improve to R² = 0.92, whereas AutoGluon was more variable across folds and stabilized at a lower value.

The significance annotations identify train–validation differences but should not automatically be interpreted as conventional sample-level overfitting. Under DOB-SCV, training and validation sets contain different dosing conditions; the observed gaps therefore combine model-fit effects with the biological difficulty of generalizing to unseen nutrient conditions. The absence of uniformly low training and validation trajectories argues against broad underfitting, but persistent gaps for several targets indicate that condition-level generalization was not fully saturated.

### 3.4 SHAP analysis identifies target-specific drivers and architecture-dependent attribution consistency

Paired Kernel SHAP beeswarm plots were used to examine how each architecture transformed the 17-feature cultivation record into mineral-specific predictions (Figs. 5 and 6). The plots retain both attribution magnitude and direction, showing whether low or high feature values shifted predicted concentrations below or above the model-specific background expectation.

**Fig. 5.**
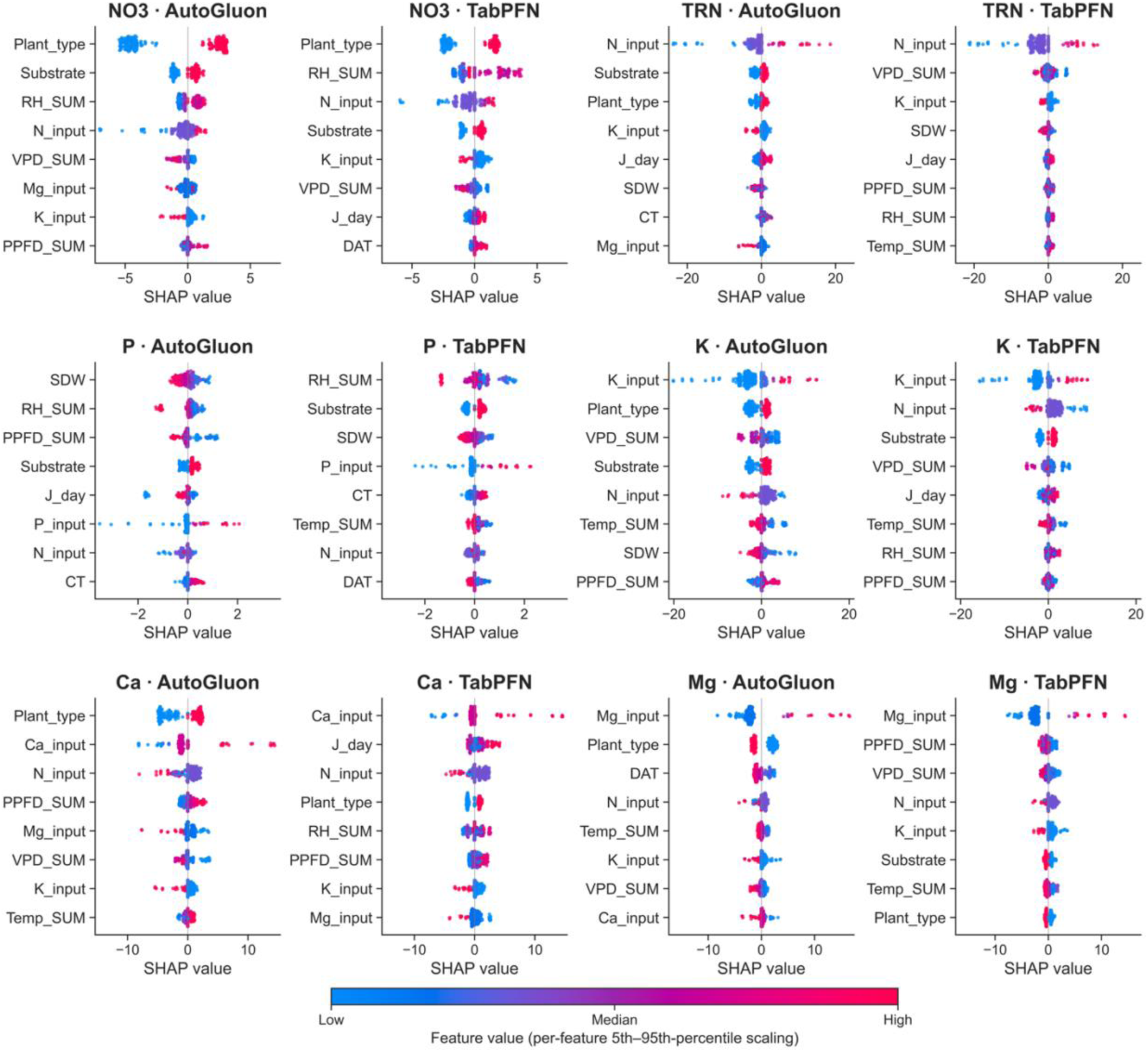
Paired Shapley additive explanations (SHAP) beeswarm summaries for the six nitrogen and macronutrient targets. Each target (NO₃⁻, total reduced nitrogen, P, K, Ca and Mg) is shown for AutoGluon 1.5 (left) and TabPFN v3 (right). Each point represents one of 250 explained instances, positioned by its SHAP value and colored by the corresponding feature value after within-feature 5th–95th percentile scaling. The eight most influential features are ordered independently for each model by mean absolute SHAP value. Paired panels share a symmetric SHAP-value axis. Model-agnostic Kernel SHAP used the first Distribution-Optimally-Balanced Stratified Cross-Validation (DOB-SCV) fold, five k-means background centroids and 2,048 coalitions; TabPFN explanations used the 32-member inference ensemble.

**Fig. 6.**
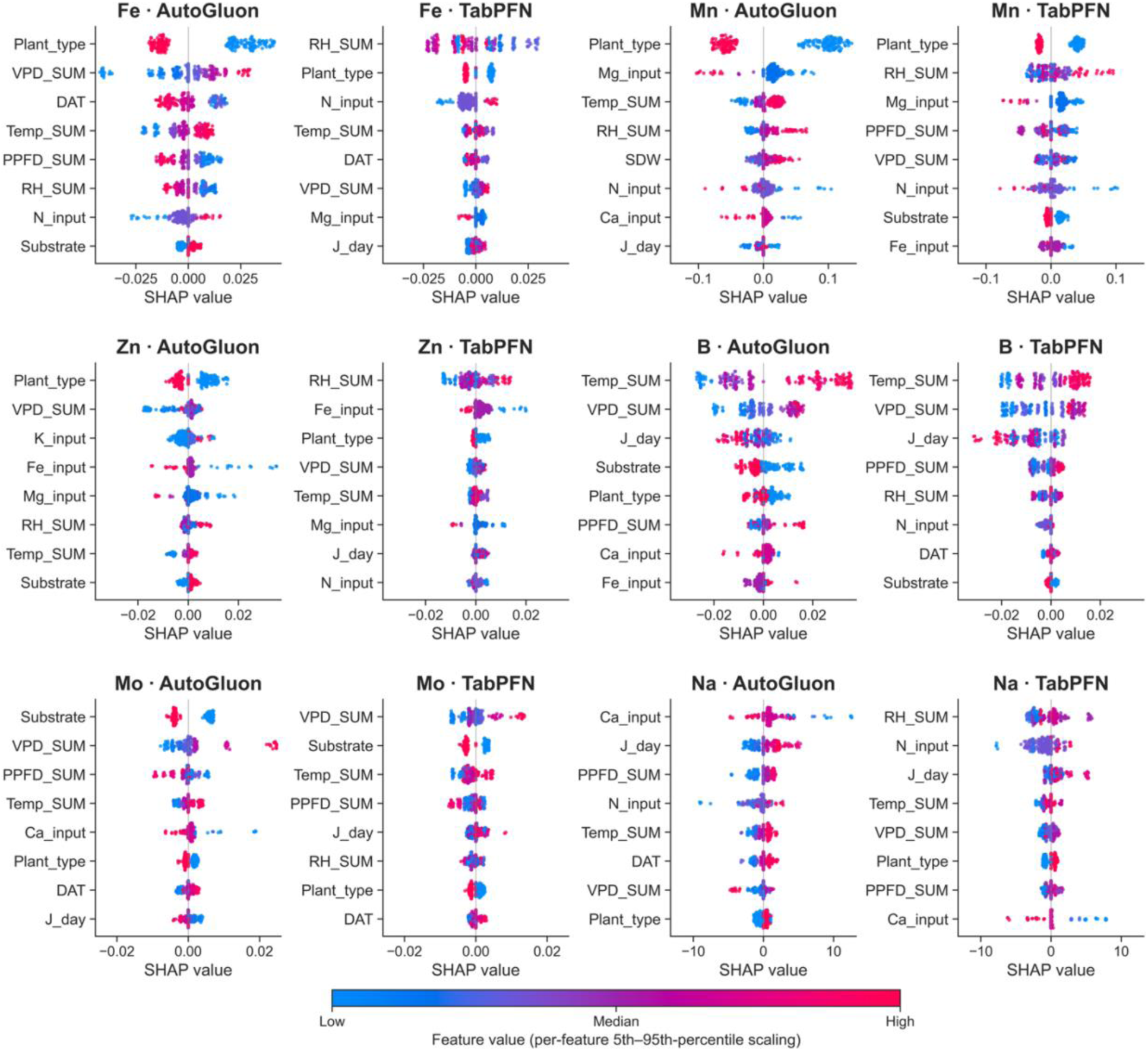
Paired Shapley additive explanations (SHAP) beeswarm summaries for the six micronutrient and low-abundance targets (Fe, Mn, Zn, B, Mo and Na). Panel pairing, point definition, within-feature 5th–95th percentile color scaling, model-specific ordering of the eight highest-ranked features, shared within-pair SHAP-value axes and the Kernel SHAP configuration are identical to those in Fig. 5.

For the nitrogen and macronutrient targets, direct mineral inputs were prominent but acted within broader physiological and environmental contexts. Plant type, substrate, RH_SUM and N_input were the principal contributors to NO₃⁻ predictions. N_input dominated TRN attribution in both architectures, with higher N supply generally shifting predicted TRN upward, while VPD_SUM, K_input, SDW and temporal variables added information. P_input contributed to P predictions, although SDW, RH_SUM, substrate and cumulative light exposure were similarly or more influential in one or both models. K_input was the leading K predictor, and Ca_input and Mg_input were prominent for their corresponding targets. Higher K_input, Ca_input and Mg_input generally produced positive SHAP shifts for the respective mineral outputs.

Micronutrient predictions relied more strongly on plant identity, environmental history and the broader nutrient background. Plant type and environmental variables repeatedly ranked highly for Fe, Mn and Zn. B was dominated by cumulative temperature, VPD and temporal descriptors in both architectures. Mo also depended strongly on substrate, VPD_SUM, PPFD_SUM, Temp_SUM and chronology, whereas Mo_input did not appear among the eight highest-ranked features. This pattern should be interpreted in the context of the single-species Mo experiment and its experiment-specific concentration gradient. Na predictions were associated with Ca_input, N_input, RH_SUM, J_day and other environmental variables. However, these attributions cannot be interpreted as independent physiological drivers because Na supply co-varied with several target gradients.

Cross-model attribution agreement was target dependent (Fig. 7). Agreement was highest for TRN (R² = 0.76), NO₃⁻ (R² = 0.75), K and Mo (R² = 0.74 each), and Mg (R² = 0.71). Intermediate agreement was observed for B (R² = 0.63), Mn (R² = 0.59) and P (R² = 0.53), whereas agreement was weaker for Ca (R² = 0.44), Fe (R² = 0.20), Na (R² = 0.17) and Zn (R² = 0.08). Comparable predictive performance therefore did not necessarily imply that the architectures used the same feature hierarchy.

**Fig. 7.**
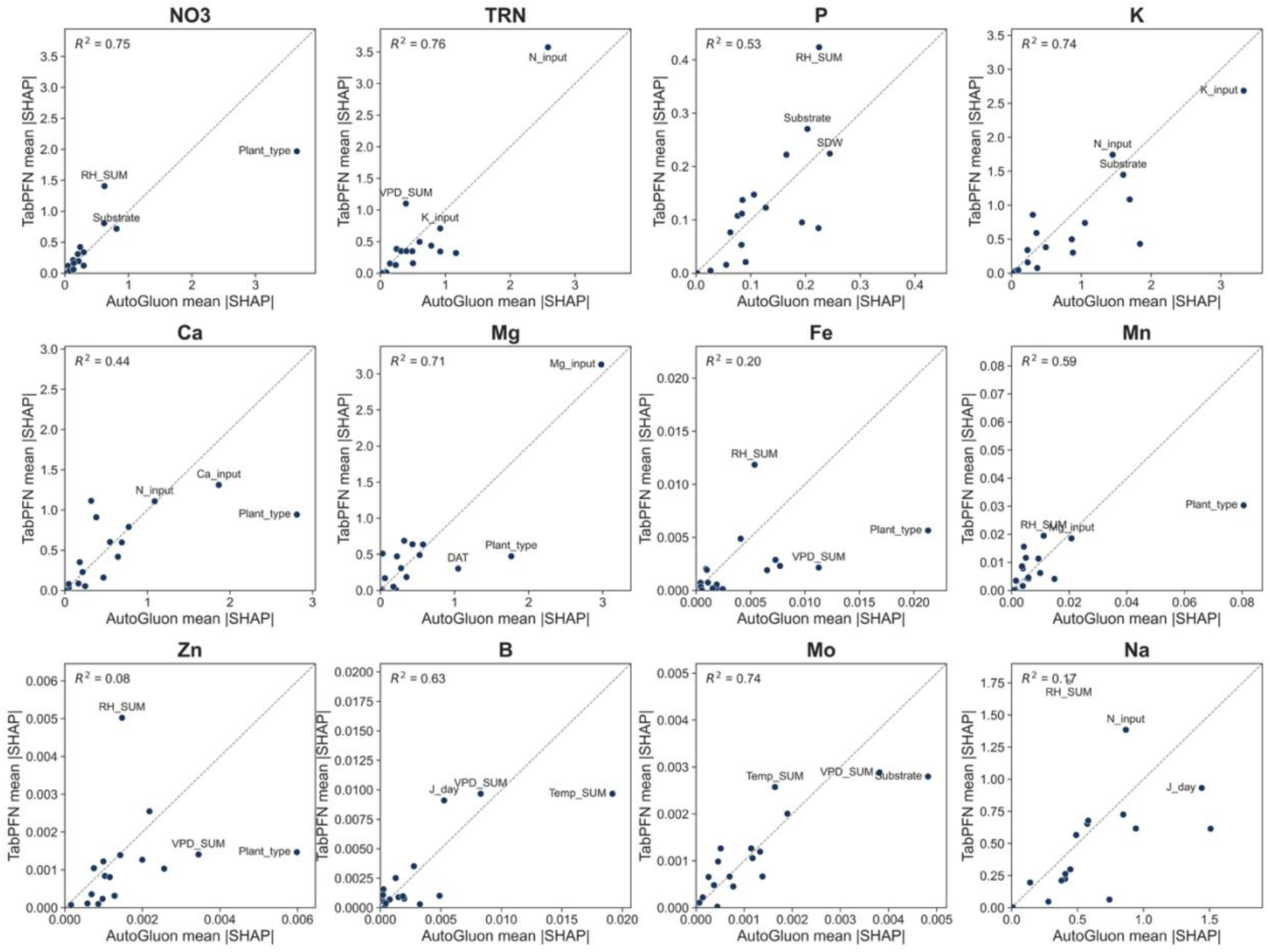
Cross-model comparison of global feature importance for the 12 harvested-leaf mineral targets. Each point represents one of the 17 input features, positioned by its mean absolute Shapley additive explanations (SHAP) value across the same 250 explained instances for AutoGluon 1.5 (x-axis) and TabPFN v3 (y-axis). inset values report squred Pearson correlation coefficients, denoted R², between the model-specific feature-importance vectors. Dashed lines indicates the 1:1 reference (y=x), representing equal attribution magnitudes.

Supplementary robustness analyses showed that SHAP feature rankings were highly stable to coalition count, background size and explained-instance selection, with mean Spearman’s rank correlation coefficients (ρ) generally exceeding 0.95 (Table S6). Fold-to-fold stability was lower but remained substantial (ρ = 0.87–0.94), as expected because each DOB-SCV fold withheld a different set of dosing conditions. Interpretability claims are therefore strongest where influential beeswarm features also show cross-model and cross-fold consistency.

### 3.5 External commercial validation reveals target-specific calibration under reduced feature availability

External transfer was evaluated with models retrained on the 15 variables available in the grower-operated cohort, with cumulative transpiration and shoot dry weight excluded from training and inference.

Because all 99 commercial plants received the same fertigation formulation and substrate, nutrient-treatment inputs contained no between-plant variation. The primary external evaluation statistic was multiplicative bias, defined as mean predicted divided by mean observed concentration, which evaluates agreement in the absolute concentration rather than explained sample-level variance (Fig. 8).

**Fig. 8.**
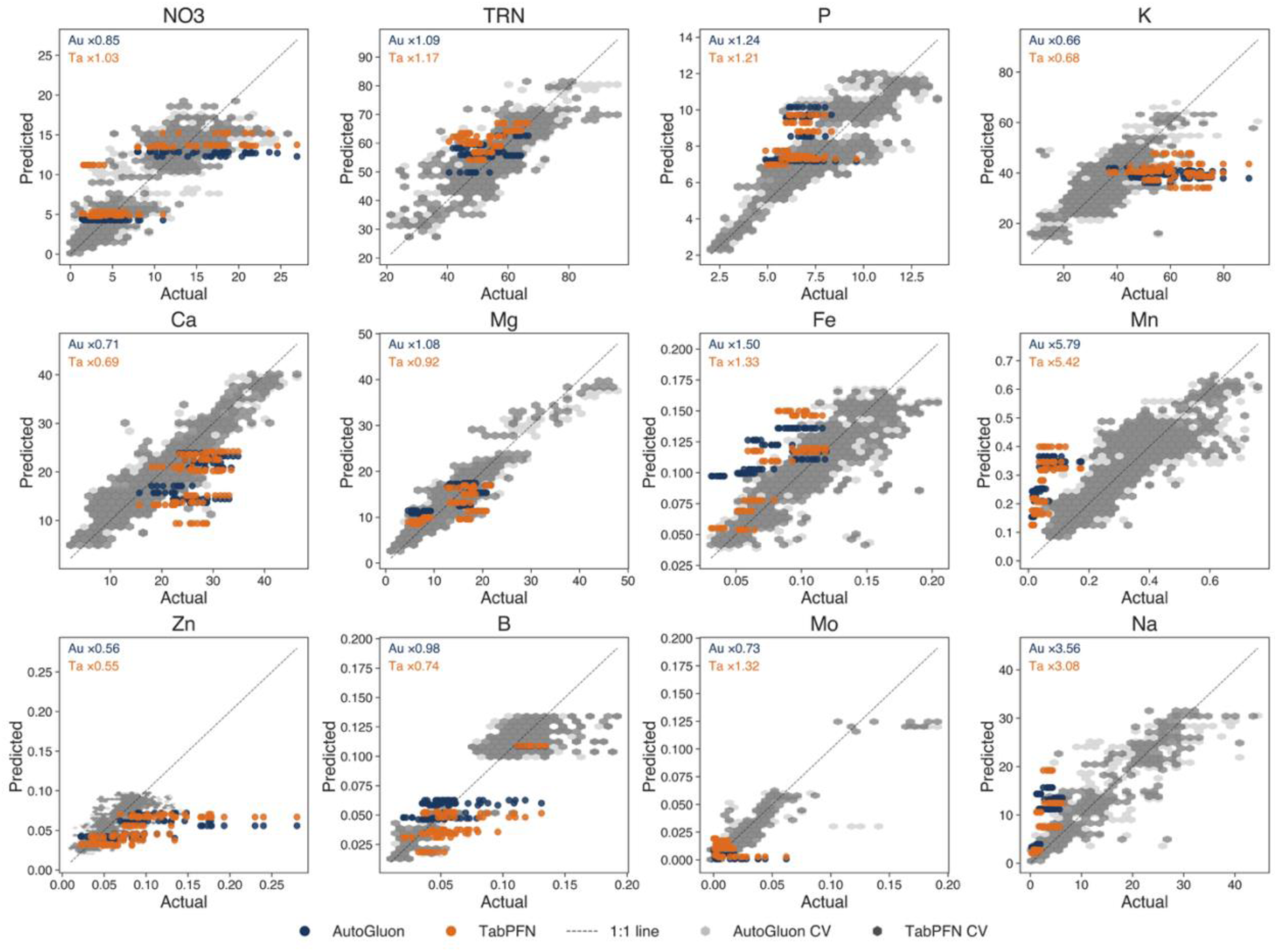
External commercial validation of reduced-feature models. Scatter plots compare analytically measured harvested-leaf concentrations (mg g⁻¹ DW) for 99 plants from nine commercial harvest cycles with predictions from AutoGluon 1.5 and TabPFN v3 models retrained without CT and SDW. Blue and orange points denote commercial predictions from AutoGluon (Au) and TabPFN (Ta), respectively; gray hexagons show the pooled internal out-of-fold prediction domain for the corresponding reduced-feature models. Insets report multiplicative bias (mean predicted/mean observed): ×1.00 is unbiased, values above 1 indicate overprediction and values below 1 indicate underprediction. Dashed lines denote 1:1 agreement.

The closest central calibration was obtained for NO₃⁻ with TabPFN (×1.03), B with AutoGluon (×0.98) and Mg with both architectures (×1.08 for AutoGluon and ×0.92 for TabPFN). TRN was moderately overestimated (×1.09 and ×1.17), and P was overestimated by 21–24% (×1.24 and ×1.21).

Several targets showed consistent directional bias across architectures. K was underestimated (×0.66 and ×0.68), as were Ca (×0.71 and ×0.69) and Zn (×0.56 and ×0.55). Fe was overestimated (×1.50 and ×1.33). The largest departures occurred for Mn (×5.79 and ×5.42) and Na (×3.56 and ×3.08), both of which were strongly overestimated. Mo calibration differed between architectures: AutoGluon underestimated the commercial mean (×0.73), whereas TabPFN overestimated it (×1.32). B also showed a model-dependent difference, with AutoGluon nearly unbiased and TabPFN underestimated the mean by 26%.

The controlled-domain out-of-fold clouds provide context for these biases but overlap with the commercial points does not itself demonstrate accurate transfer. Because nutrient inputs were fixed within the commercial cohort, predictions frequently formed compressed bands driven mainly by plant type, temporal descriptors and microclimate. The shared direction of AutoGluon and TabPFN bias for K, Ca, Fe, Mn, Zn and Na suggests that these failures were primarily associated with the deployment domain and missing feature information rather than with one architecture.

## 4. Discussion

### 4.1 Predictive ionomics remains robust when validation is restricted to unseen nutrient conditions

The central finding of this study is that the harvested-leaf ionome of leafy vegetables contains substantial, learnable predictive structure. Under leakage-free grouped cross-validation, AutoGluon and TabPFN predicted 12 mineral targets with pooled out-of-fold R² values of 0.72–0.92. These estimates were obtained by withholding complete dosing conditions rather than individual biological replicates; the models were therefore required to predict plants grown under nutrient conditions absent from the corresponding training set. This distinction is fundamental because conventional random cross-validation can yield optimistic estimates when related or hierarchically structured observations are divided between training and validation sets. Validation design should therefore reflect both the dependence structure of the data and the intended prediction task (Moreno-Torres et al., 2012; Roberts et al., 2017).

The similarity in overall performance between AutoGluon and TabPFN further strengthens this conclusion. AutoGluon constructs task-specific stacked ensembles from heterogeneous base learners, whereas TabPFN applies a pretrained prior-data-fitted network to tabular prediction. Neither architecture consistently dominated across the ionomic panel. B, Mg and Ca were among the most accurately predicted targets in both frameworks, whereas TRN, K, Mn and Zn remained more challenging. Mo was an informative exception, with TabPFN substantially outperforming AutoGluon. The overall predictive signal therefore appears to reside primarily in the experimental feature–phenotype structure, while architecture-specific inductive biases remain relevant for individual targets.

The scope of this generalization must nevertheless be defined precisely. DOB-SCV evaluates transfer to unseen nutrient-dose conditions while retaining representation of all 16 cultivation experiments in every fold. It is therefore more stringent than replicate-level random partitioning, but it is not equivalent to withholding an entire experiment, species, season or production system. Structured cross-validation is most informative when its grouping structure corresponds to the intended deployment question (Roberts et al., 2017). Here, internal validation asks whether the models can interpolate across previously unseen nutrient conditions within the broader experimental landscape, whereas the independent commercial cohort separately tests transfer beyond that landscape.

### 4.2 The leaf ionome is predictable because it is structured, but not because its elements act independently

Ionomics is founded on the premise that elemental composition represents an integrated phenotype rather than a collection of independent nutrient measurements (Salt et al., 2008). Subsequent work has reinforced this systems-level view, showing that ionomic profiles encode genetic, environmental and physiological information and that combinations of elements may be more informative than individual traits considered in isolation (Baxter, 2015; Huang and Salt, 2016). Our results extend this concept to predictive modeling of edible leafy vegetables.

Targeted nutrient gradients generated broad, non-Gaussian and species-dependent distributions, and perturbations frequently coincided with shifts in mineral outputs other than the directly manipulated element. Such coordinated responses are biologically plausible because nutrient acquisition and homeostasis are interconnected through transport, signaling, ionic balance and metabolic regulation (de Bang et al., 2021; Hanikenne et al., 2021). The strong contribution of plant type in both the KDE and SHAP analyses is also consistent with evidence that ionomic composition carries substantial genetic and taxonomic structure and interacts strongly with the growth environment (Huang and Salt, 2016; Stein et al., 2017).

The experimental design also imposes an important boundary on this interpretation. Na and S partly co-varied with specific target gradients because they were introduced as counter-ions during nutrient-solution formulation (Table S1). Their off-target responses therefore cannot be interpreted in the same way as those of elements whose supplied concentrations remained constant. The strongest evidence for emergent multi-ionomic responses comes from off-target changes in K, Ca, Mg and micronutrients whose external supply was controlled within the corresponding gradient. In contrast, leaf Na reflects a mixture of plant regulation and engineered fertigation co-variation.

This distinction is particularly relevant to the strong internal predictability of Na. Within the controlled design, nutrient inputs contain information about the counter-ion structure associated with treatment identity, enabling the models to reconstruct Na concentrations accurately. That experimental co-variation was absent from the fixed commercial formulation, where Na showed substantial calibration bias. We therefore interpret Na as a highly predictable but partially design-confounded target in this dataset, not as strong evidence of emergent physiological crosstalk. Separating engineered from biological covariance is essential if predictive ionomics is to progress from statistical mapping toward mechanistic inference.

### 4.3 Physiological and environmental histories provide information that nutrient supply alone cannot capture

A second major finding is that harvested-leaf mineral concentration was not predicted solely from fertilizer composition. Across targets, the models repeatedly used plant type, substrate, cumulative environmental exposure, cumulative transpiration, shoot dry weight and developmental descriptors together with nutrient inputs. This is physiologically reasonable because the concentration measured at harvest is the endpoint of processes operating throughout the cultivation cycle: nutrient acquisition, water transport, allocation, metabolism and biomass accumulation.

The role of water relations is particularly clear for elements associated with xylem transport. Calcium is a well-established example because its delivery to aerial organs is closely coupled to water flow and transpiration, although subsequent cellular transport and storage also influence tissue distribution (Gilliham et al., 2011; White and Broadley, 2003). Atmospheric vapor pressure deficit similarly regulates stomatal behavior and whole-plant water loss, linking environmental demand to transpiration and growth over time (Grossiord et al., 2020). These relationships provide a plausible basis for the repeated contribution of CT, VPD_SUM and RH_SUM to mineral prediction.

Biomass provides a complementary mechanism. Tissue concentration depends not only on the amount of mineral acquired but also on the biomass over which that mineral is distributed. Rapid dry-matter accumulation can therefore reduce measured nutrient concentration even when total nutrient acquisition increases—the classical growth-dilution effect (Jarrell and Beverly, 1981). The recurrent contribution of SDW in the SHAP analyses is consistent with explicitly representing this dilution component.

The distinction between NO₃⁻ and TRN further illustrates the value of physiological context. Leaf nitrate reflects the balance among nitrate uptake, transport, storage and reduction and is strongly influenced by nutrient supply and environmental conditions in leafy vegetables (Bian et al., 2020). TRN, in contrast, integrates reduced nitrogen incorporated into metabolically and structurally diverse pools. Under DOB-SCV, NO₃⁻ was predicted more accurately than TRN, suggesting that the measured nutrient and environmental descriptors captured nitrate accumulation more completely than downstream reduced-N metabolism.

Cumulative environmental variables should nevertheless be interpreted as predictive descriptors rather than as direct demonstrations of mechanism. One experimental cycle required xLSTM reconstruction of missing internal microclimate telemetry, and reconstruction accuracy differed among variables. VPD showed good point-wise performance (R² = 0.813), substantially exceeding PPFD (R² = 0.621; Fig. S4). After temporal integration, Temp_SUM and RH_SUM showed minimal cumulative bias, whereas PPFD_SUM and VPD_SUM retained whole-window biases of +12.6% and +6.6%, respectively (Fig. S5). The VPD result therefore reflects the accumulation of modest systematic error during integration rather than poor reconstruction of its temporal dynamics. Removing the imputed cultivation cycle changed ionomic prediction by only |ΔR²| = 0.013 on average, indicating that the central results were insensitive to inclusion of that experiment. SHAP effects involving reconstructed cumulative variables should accordingly be interpreted in light of this quantified uncertainty.

### 4.4 Learning curves reveal where additional biological information is most valuable

Learning curves provide more than a diagnostic of model fit: they describe how generalization changes with the amount of training information and can indicate whether additional data are likely to improve prediction (Viering and Loog, 2023). This is particularly relevant in biological ML, where datasets are costly to generate and may occupy highly structured regions of phenotype space (Mey et al., 2021; van Dijk et al., 2021). In ionomics, each new observation requires destructive harvest, sample preparation and chemical analysis, making the allocation of additional experiments a practical scientific question.

The grouped learning curves reveal an important distinction from replicate-level validation. Several targets acquired most of their predictive performance at relatively small training fractions, but this did not always produce convergence between training and validation trajectories. B showed the clearest convergence, whereas NO₃⁻, P, Ca, Mg and Na reached comparatively high validation performance while retaining modest condition-level gaps. TRN, K, Mn and Zn showed more persistent separation, indicating greater sensitivity to the coverage of nutrient conditions.

Under DOB-SCV, these gaps should not automatically be labeled conventional overfitting. Every validation fold comprises complete dosing conditions absent from training; train–validation separation therefore combines model-fitting effects with the biological difficulty of interpolating across unobserved regions of the nutrient-gradient design. This is precisely why validation structure matters: a model can fit observed conditions well yet remain less certain when it encounters a new nutrient combination or concentration level (Roberts et al., 2017).

This interpretation also changes what ‘more data’ should mean in future experiments. Additional biological replicates within densely sampled dosing conditions may provide less generalization benefit than greater diversity of training conditions. Our results therefore support prioritizing new intermediate and boundary nutrient levels, additional nutrient combinations, species–substrate combinations and independent cultivation periods, particularly for TRN, K, Mn and Zn. The learning curves thus become an experimental-design tool by identifying where new destructive ionomic measurements are most likely to expand the useful prediction domain.

### 4.5 SHAP reveals biologically coherent predictive structure, but explanation is not mechanism

SHAP provided a complementary view of the predictive models by identifying the variables that contributed most strongly to individual predictions. SHAP assigns local feature contributions relative to a model-specific baseline within a game-theoretic framework. Several macronutrient targets showed intuitive direct-input relationships: N_input was prominent for TRN, K_input for K, Ca_input for Ca and Mg_input for Mg. These inputs nevertheless operated within broader feature structures containing plant identity, substrate, environmental integrals, biomass and temporal descriptors. This distributed attribution is consistent with the systems-level nature of plant mineral nutrition rather than with simple fertilizer-to-tissue transfer functions.

Cross-model comparison adds an important layer to this interpretation. Agreement was strongest for TRN, NO₃⁻, K, Mo and Mg, for which the two architectures recovered relatively similar global feature hierarchies. Agreement was intermediate for B, Mn and P and weaker for Ca, Fe, Na and particularly Zn. Predictive and explanatory agreement are therefore not equivalent: two architectures can achieve similar out-of-fold accuracy while using different combinations of correlated predictors.

This distinction deserves particular attention in structured biological datasets. Kernel SHAP explanations can be sensitive to dependencies among input variables because correlated features can share or redistribute attribution in ways that complicate direct mechanistic interpretation (Aas et al., 2021). More generally, post hoc explanations of complex predictive models should not substitute for experimentally identifiable causal mechanisms (Rudin, 2019) . In this study, nutrient inputs, plant type, substrate, seasonality and cumulative environmental variables are not statistically independent, making causal language inappropriate.

Na provides a clear example. Its weak cross-model attribution agreement, together with known counterion co-variation in the nutrient formulations, means that RH_SUM, N_input or Ca_input should be interpreted as variables used to reconstruct Na predictions within this design, not as evidence that they independently regulate Na accumulation. Similar caution applies to Mo, for which only one plant type was represented in the gradient experiment and Mo_input was not among the leading global SHAP features.

The supplementary robustness analysis nevertheless showed that the SHAP procedure itself was stable. Feature rankings were highly conserved across coalition counts, background sizes and explained-instance samples, with mean Spearman’s rank correlation coefficients generally > 0.95. Fold-to-fold agreement also remained substantial despite each fold withholding different dosing conditions (Table S6). The main uncertainty therefore lies less in numerical SHAP convergence than in the biological interpretation of correlated predictors.

The strongest explanatory evidence arises where four lines of support coincide: high leakage-free predictive performance, strong within-model SHAP importance, agreement between AutoGluon and TabPFN, and consistency with established plant physiology. Under this framework, SHAP is most useful as a hypothesis-generating tool that prioritizes nutrient–environment–physiology relationships for controlled testing, not as proof of causal regulation.

### 4.6 Commercial validation exposes calibration limits that internal cross-validation cannot reveal

Even rigorous internal cross-validation cannot establish whether a model will perform in a different production system. Predictive models should therefore be evaluated outside their development domain when the intended application involves new populations or operating conditions (Ramspek et al., 2020; Steyerberg and Harrell, 2016). The grower-operated cohort served this purpose and introduced several simultaneous changes: commercial fertigation, high planting density, a shared root zone and the absence of plant-level CT and SDW measurements.

This external test also differs statistically from DOB-SCV. All 99 commercial plants received the same fertilizer formulation and substrate, so most nutrient-input variables carried no between-plant information. The reduced models varied predictions primarily through plant identity, developmental timing and microclimatic history. Under these conditions, the first question is not how much within-cohort variance is explained but whether predictions are calibrated in absolute concentration. Calibration is distinct from discrimination and can deteriorate when a model is transferred to a population or domain that differs from its development data (Van Calster et al., 2019).

The multiplicative-bias analysis revealed a clear target-specific hierarchy. The closest calibration was observed for TabPFN NO₃⁻, AutoGluon B and both Mg models, whose predicted means approached the observed commercial concentrations. TRN showed moderate overestimation, and P was overestimated by approximately one fifth. Both architectures underestimated K, Ca and Zn and overestimated Fe. The largest calibration failures occurred for Mn and Na, whose predicted concentrations substantially exceeded the commercial observations.

The shared direction of many errors is informative. AutoGluon and TabPFN use different computational strategies, yet both underpredicted K, Ca and Zn and overestimated Fe, Mn and Na. The dominant limitation therefore appears to lie in the deployment-domain information rather than in a failure unique to one architecture. Dataset shift can alter predictor–outcome relationships when models move between environments, making external validation essential even after strong internal results (Moreno-Torres et al., 2012; Steyerberg and Harrell, 2016).

The particularly poor external calibration of Na reinforces the counter-ion limitation. A relationship that was strongly predictable within the controlled nutrient-gradient design did not transfer to the fixed commercial formulation, consistent with part of the internal signal being specific to the experimental solution structure. This illustrates why predictive accuracy alone cannot establish biological generality.

The controlled-domain prediction clouds behind the commercial observations remain useful as a visual reference for the numerical range learned by the models, but overlap with that envelope should not be interpreted as successful external validation. A commercial prediction can remain within the internal range and still be systematically biased. Absolute calibration against measurements from the intended production system is therefore more informative for deployment than visual overlap alone.

### 4.7 From predictive ionomics to deployable crop monitoring

The broader value of predictive ionomics lies in connecting nutrient management with measurable plant and environmental state. Precision agriculture seeks to replace uniform management with decisions informed by spatial, temporal and biological variability (Gebbers and Adamchuk, 2010), while smart-farming frameworks increasingly integrate sensors, computational models and management data into continuous decision-support systems (Wolfert et al., 2017). ML provides an important analytical layer within this transition, but agricultural value depends on performance under realistic operating conditions rather than on benchmark accuracy alone (Kamilaris and Prenafeta-Boldú, 2018; van Dijk et al., 2021).

Our results suggest a staged route toward deployment. Sensor-rich research and advanced controlled-environment systems can use the full feature set, including cumulative transpiration and biomass, to maximize predictive resolution. A grower-deployable model can instead operate with the 15 features available from fertigation records, crop descriptors and environmental monitoring. The external analysis indicates that this reduced representation may provide useful absolute estimates for selected targets, particularly NO₃⁻ and Mg, but should not be assumed to generalize equally across the ionome.

A third, potentially more practical pathway is local calibration. A small number of analytically measured commercial reference samples could determine whether target-specific predictions require intercept or scale correction before use. Model updating and recalibration after external validation are established approaches when predictive relationships remain informative but absolute calibration shifts between development and deployment populations (Van Calster et al., 2019; van Dijk et al., 2021). For targets showing severe compression or large bias, however, calibration alone may be insufficient and additional explanatory features may be required.

Future validation should also broaden the commercial domain. The present cohort includes repeated harvest cycles but only one commercial fertilizer formulation and production system. Multi-farm evaluation across fertilizer recipes, planting densities, seasons, substrates and crop types will be required before broad field transferability can be claimed. This expansion should retain grouped validation so that evaluation units represent genuinely unseen management conditions rather than additional biological replicates from conditions already encountered during training.

The practical endpoint should therefore not be a universal ‘ionome model’ but a target-aware monitoring system in which every mineral has its own validated range, calibration status and uncertainty. A model that is reliable for NO₃⁻ or Mg should not automatically be considered reliable for Mn or Na. Explicit target-specific validation is essential if predictive ionomics is to become a credible component of precision-fertilization decision support.

### 4.8 Conclusion

This study demonstrates that designed nutrient-gradient phenotyping, combined with continuous plant– environment monitoring and modern tabular learning, predicts a broad harvested-leaf ionome. Under leakage-free validation of complete unseen dosing conditions, AutoGluon and TabPFN retained substantial predictive performance across 12 targets, while learning curves identified mineral-specific differences in data sufficiency and generalization. SHAP revealed interpretable combinations of nutrient, plant, substrate, physiological and environmental predictors, but cross-model comparisons also showed that explanatory stability is target dependent and should not be equated with causal mechanism.

The independent commercial cohort provides an equally important boundary to these findings. Some targets retained useful absolute calibration after physiological variables were removed, whereas others showed substantial systematic bias. Predictive ionomics is therefore technically feasible, but deployment remains a mineral-specific calibration and sensing problem. By combining designed perturbation, leakage-aware validation, physiological phenotyping, explainable modeling and independent commercial testing, this framework provides a rigorous foundation for future data-driven tools for crop mineral management and nutritional-quality monitoring.

## CRediT authorship contribution statement

Itamar Shenhar: Data curation, Formal analysis, Investigation, Methodology, Software, Visualization, Validation, Writing - original draft. Miguel R. Pebes-Trujillo: Investigation, Writing - review & editing. Aravind Harikumar: Investigation, Writing - review & editing. Zhitong Zhao: Investigation, Writing - review & editing. Li Yi Tan: Investigation, Writing - review & editing. Magdiel Inggrid Setyawati: Investigation, Writing - review & editing. Jie He: Resources, Writing - review & editing. Ittai Herrmann: Writing - review & editing. Kee Woei Ng: Project administration, Funding acquisition, Writing - review & editing. Matan Gavish: Funding acquisition, Writing - review & editing. Menachem Moshelion: Conceptualization, Methodology, Project administration, Supervision, Writing - review & editing.

## Data availability

The data supporting the findings of this study are available from the corresponding author upon reasonable request. The complete analysis pipeline, including the Slurm scripts used for leakage-free cross-validated training, SHAP attribution and external commercial-cohort validation is openly available under the MIT license at: https://github.com/itamarshenhar/predictive-ionomics.

## Declaration of generative AI and AI-assisted technologies in the writing process

During the preparation of this work the authors used ChatGPT (OpenAI) to improve the readability and language and Claude (Anthropic) to develop the code to support the study. After using these tools, the authors reviewed and edited the content as needed and take full responsibility for the content of the published article.

## Acknowledgments

The Israel–Singapore Urban Farm (iSURF) is supported by the National Research Foundation, Prime Minister’s Office, Singapore, under the Campus for Research Excellence and Technological Enterprise (CREATE) programme [NRF2020-THE003-0008]. This work was also supported in part by the Hebrew University of Jerusalem Center for Interdisciplinary Data Science Research.

We gratefully acknowledge assistance from iSURF collaborators at Netatech Pte Ltd, Singapore, Mr. David Tan and the Singapore Food Agency. We also thank Dr. Qin Lin for her assistance with the development and implementation of the laboratory analytical procedures and Mrs. Maria Carmencita Morales for administrative support.

## References

1. Aas, K., Jullum, M., Løland, A., 2021. Explaining individual predictions when features are dependent: More accurate approximations to Shapley values. Artif. Intell. 298, 103502. 10.1016/J.ARTINT.2021.103502

2. Allen, S.E., 1989. Chemical analysis of ecological materials, in: Allen, D. (Ed.), Chemical Analysis of Ecological Materials. Blackwell Scientific, Oxford, pp. 46–61.

3. Baxter, I., 2015. Should we treat the ionome as a combination of individual elements, or should we be deriving novel combined traits? J. Exp. Bot. 66, 2127–2131. 10.1093/JXB/ERV040

4. Bian, Z., Wang, Y., Zhang, X., Li, T., Grundy, S., Yang, Q., Cheng, R., 2020. A Review of Environment Effects on Nitrate Accumulation in Leafy Vegetables Grown in Controlled Environments. Foods 9, 732. 10.3390/FOODS9060732

5. Bonhomme, R., 2000. Bases and limits to using ‘degree.day’ units. European Journal of Agronomy 13, 1–10. 10.1016/S1161-0301(00)00058-7

6. Dalal, A., Shenhar, I., Bourstein, R., Mayo, A., Grunwald, Y., Averbuch, N., Attia, Z., Wallach, R., Moshelion, M., 2020. A Telemetric, Gravimetric Platform for Real-Time Physiological Phenotyping of Plant–Environment Interactions. JoVE (Journal of Visualized Experiments) 2020, e61280. 10.3791/61280

7. de Bang, T.C., Husted, S., Laursen, K.H., Persson, D.P., Schjoerring, J.K., 2021. The molecular– physiological functions of mineral macronutrients and their consequences for deficiency symptoms in plants. New Phytologist 229, 2446–2469. 10.1111/NPH.17074

8. Erickson, N., Mueller, J., Shirkov, A., Zhang, H., Larroy, P., Li, M., Smola, A., 2020. AutoGluon-Tabular: Robust and Accurate AutoML for Structured Data. ArXiv. 10.48550/ARXIV.2003.06505

9. Fan, X., Zhou, X., Chen, H., Tang, M., Xie, X., 2021. Cross-Talks Between Macro- and Micronutrient Uptake and Signaling in Plants. Front. Plant Sci. 12, 663477. 10.3389/FPLS.2021.663477

10. Faust, J.E., Logan, J., 2018. Daily Light Integral: A Research Review and High-resolution Maps of the United States. HortScience 53, 1250–1257. 10.21273/HORTSCI13144-18

11. Gebbers, R., Adamchuk, V.I., 2010. Precision Agriculture and Food Security. Science (1979). 10.1126/SCIENCE.1183899

12. Gilliham, M., Dayod, M., Hocking, B.J., Xu, B., Conn, S.J., Kaiser, B.N., Leigh, R.A., Tyerman, S.D., 2011. Calcium delivery and storage in plant leaves: exploring the link with water flow. J. Exp. Bot. 62, 2233–2250. 10.1093/JXB/ERR111

13. Grossiord, C., Buckley, T.N., Cernusak, L.A., Novick, K.A., Poulter, B., Siegwolf, R.T.W., Sperry, J.S., McDowell, N.G., 2020. Plant responses to rising vapor pressure deficit. New Phytologist 226, 1550–1566. 10.1111/NPH.16485

14. Halperin, O., Gebremedhin, A., Wallach, R., Moshelion, M., 2017. High-throughput physiological phenotyping and screening system for the characterization of plant–environment interactions. The Plant Journal 89, 839–850. 10.1111/TPJ.13425

15. Hanikenne, M., Esteves, S.M., Fanara, S., Rouached, H., 2021. Coordinated homeostasis of essential mineral nutrients: a focus on iron. J. Exp. Bot. 72, 2136–2153. 10.1093/JXB/ERAA483

16. He, J., Chua, E.L., Qin, L., 2020. Drought does not induce crassulacean acid metabolism (CAM) but regulates photosynthesis and enhances nutritional quality of Mesembryanthemum crystallinum. PLoS One 15, e0229897. 10.1371/JOURNAL.PONE.0229897

17. Hollmann, N., Müller, S., Purucker, L., Krishnakumar, A., Körfer, M., Hoo, S. Bin, Schirrmeister, R.T., Hutter, F., 2025. Accurate predictions on small data with a tabular foundation model. Nature 2025 637:8045 637, 319–326. 10.1038/s41586-024-08328-6

18. Huang, X.Y., Salt, D.E.E., 2016. Plant Ionomics: From Elemental Profiling to Environmental Adaptation. Mol. Plant 9, 787–797. 10.1016/j.molp.2016.05.003

19. Jarrell, W.M., Beverly, R.B., 1981. The Dilution Effect in Plant Nutrition Studies. Advances in Agronomy 34, 197–224. 10.1016/S0065-2113(08)60887-1

20. Jones, H.G., 2013. Plants and Microclimate: A Quantitative Approach to Environmental Plant Physiology, Plants and Microclimate: A Quantitative Approach to Environmental Plant Physiology. Cambridge University Press. 10.1017/CBO9780511845727

21. Kamilaris, A., Prenafeta-Boldú, F.X., 2018. Deep learning in agriculture: A survey. Comput. Electron. Agric. 147, 70–90. 10.1016/J.COMPAG.2018.02.016

22. Kohavi, R., 1995. A Study of Cross-Validation and Bootstrap for Accuracy Estimation and Model Selection. Proceedings of the 14th International Joint Conference on Artificial Intelligence (IJCAI ‘95) 1137–1145.

23. Lundberg, S.M., Lee, S.I., 2017. A Unified Approach to Interpreting Model Predictions. Adv. Neural Inf. Process. Syst. 2017-December, 4766–4775. 10.48550/arXiv.1705.07874

24. McMaster, G.S., Wilhelm, W.W., 1997. Growing degree-days: one equation, two interpretations. Agric. For. Meteorol. 87, 291–300. 10.1016/S0168-1923(97)00027-0

25. Mey, F., Clauwaert, J., Van Huffel, K., Waegeman, W., De Mey, M., 2021. Improving the performance of machine learning models for biotechnology: The quest for deus ex machina. Biotechnol. Adv. 53, 107858. 10.1016/J.BIOTECHADV.2021.107858

26. Moreno-Torres, J.G., Saez, J.A., Herrera, F., 2012. Study on the impact of partition-induced dataset shift on k-fold cross-validation. IEEE Trans. Neural Netw. Learn. Syst. 23, 1304–1312. 10.1109/TNNLS.2012.2199516

27. National Environment Agency, 2024. Air Temperature/Rainfall/Relative Humidity/Wind Speed across Singapore (2024-2025) [WWW Document]. URL data.gov.sg (accessed 5.24.26).

28. Pedregosa, F., Varoquaux, G., Gramfort, A., Michel, V., Thirion, B., Grisel, O., Blondel, M., Prettenhofer, P., Weiss, R., Dubourg, V., 2011. Scikit-learn: Machine Learning in Python. Journal of Machine Learning Research 12, 2825–2830. 10.34894/VQ1DJA

29. Ramspek, C.L., Jager, K.J., Dekker, F.W., Zoccali, C., Van DIepen, M., 2020. External validation of prognostic models: what, why, how, when and where? Clin. Kidney J. 14, 49. 10.1093/CKJ/SFAA188

30. Roberts, D.R., Bahn, V., Ciuti, S., Boyce, M.S., Elith, J., Guillera-Arroita, G., Hauenstein, S., Lahoz-Monfort, J.J., Schröder, B., Thuiller, W., Warton, D.I., Wintle, B.A., Hartig, F., Dormann, C.F., 2017. Cross-validation strategies for data with temporal, spatial, hierarchical, or phylogenetic structure. Ecography 40, 913–929. 10.1111/ECOG.02881

31. Rudin, C., 2019. Stop explaining black box machine learning models for high stakes decisions and use interpretable models instead. Nature Machine Intelligence 2019 1:5 1, 206–215. 10.1038/s42256-019-0048-x

32. Salt, D.E., Baxter, I., Lahner, B., 2008. Ionomics and the study of the plant ionome. Annu. Rev. Plant Biol. 59, 709–733. 10.1146/ANNUREV.ARPLANT.59.032607.092942

33. Stein, R.J., Höreth, S., de Melo, J.R.F., Syllwasschy, L., Lee, G., Garbin, M.L., Clemens, S., Krämer, U., 2017. Relationships between soil and leaf mineral composition are element-specific, environment-dependent and geographically structured in the emerging model Arabidopsis halleri. New Phytologist 213, 1274–1286. 10.1111/NPH.14219

34. Steyerberg, E.W., Harrell, F.E., 2016. Prediction models need appropriate internal, internal–external, and external validation. J. Clin. Epidemiol. 69, 245–247. 10.1016/J.JCLINEPI.2015.04.005

35. Van Calster, B., McLernon, D.J., Van Smeden, M., Wynants, L., Steyerberg, E.W., Bossuyt, P., Collins, G.S., MacAskill, P., Moons, K.G.M., Vickers, A.J., 2019. Calibration: The Achilles heel of predictive analytics. BMC Med. 17, 230-. 10.1186/S12916-019-1466-7

36. van Dijk, A.D.J., Kootstra, G., Kruijer, W., de Ridder, D., 2021. Machine learning in plant science and plant breeding. iScience 24, 101890. 10.1016/j.isci.2020.101890

37. Viering, T., Loog, M., 2023. The Shape of Learning Curves: A Review. IEEE Trans. Pattern Anal. Mach. Intell. 45, 7799–7819. 10.1109/TPAMI.2022.3220744

38. White, P.J., Broadley, M.R., 2003. Calcium in Plants. Ann. Bot. 92, 487–511. 10.1093/AOB/MCG164

39. Wolfert, S., Ge, L., Verdouw, C., Bogaardt, M.J., 2017. Big Data in Smart Farming – A review. Agric. Syst. 153, 69–80. 10.1016/J.AGSY.2017.01.023

40. Zeng, X., Martinez, T.R., 2000. Distribution-balanced stratified cross-validation for accuracy estimation. Journal of Experimental & Theoretical Artificial Intelligence 12, 1–12. 10.1080/095281300146272

41. Zippenfenig, P., 2023. Open-Meteo.com Weather API. 10.5281/zenodo.7970649

