## Supplementary information for "Designed nutrient-gradient phenotyping enables predictive ionomics of tropical leafy greens"

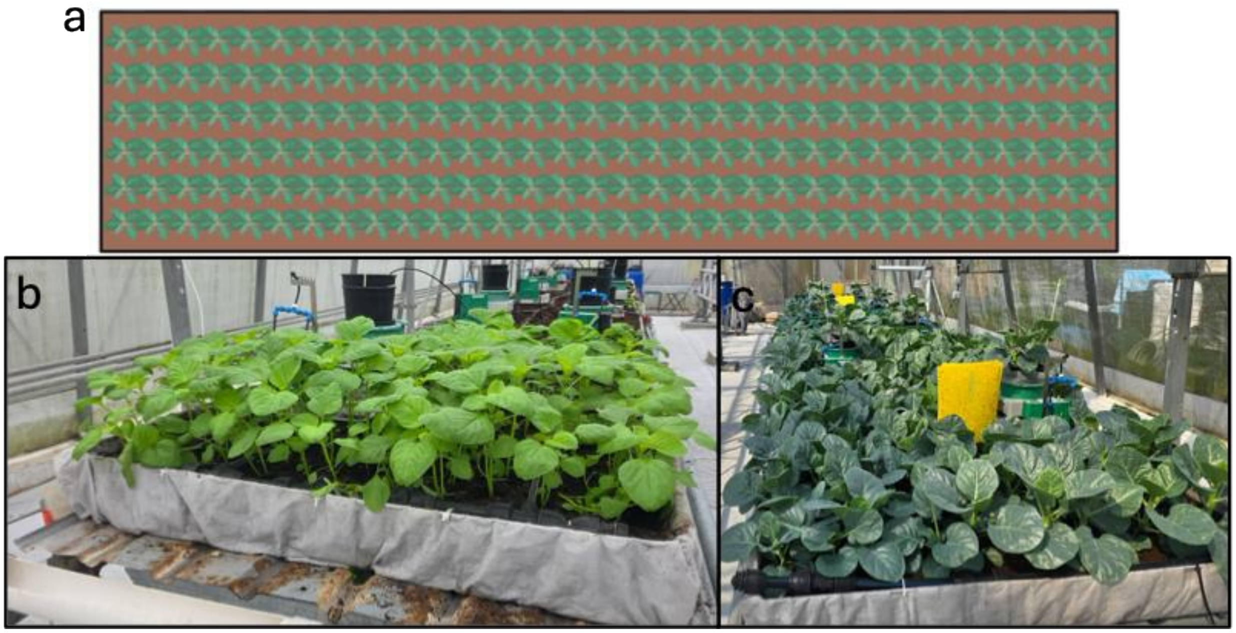


**Fig. S1.** Configuration of the grower-operated commercial side tables used for external model validation. (A) Schematic of the high-density, multirow planting arrangement used to evaluate model transfer under commercial production conditions, including canopy shading and root-zone competition within a shared substrate bed. (B) Representative photograph of Chinese spinach cultivated at high density on the side-table validation platform. (C) Representative photograph of Chinese broccoli cultivated concurrently under the same commercial production conditions. This independent system used a shared root-zone matrix and commercial fertilizer management to evaluate external transfer and calibration of the AutoGluon and TabPFN pipelines.

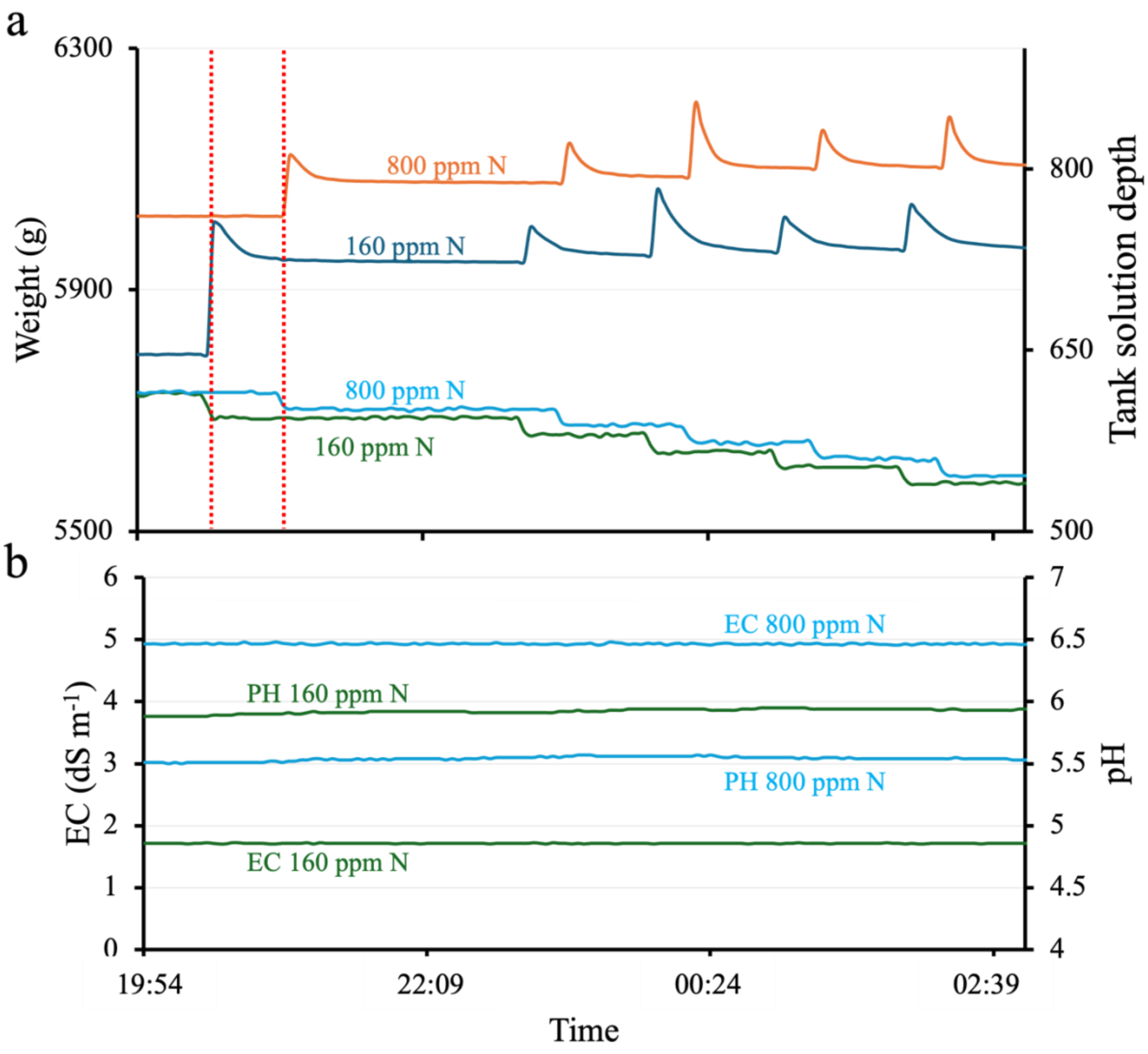


**Fig. S2.** Real-time telemetry of automated nocturnal fertigation dynamics and solution chemical stability. High-resolution time-series data were recorded at 3-min intervals by the central PlantArray data-acquisition system to monitor delivery precision. (a) Synchronized records of whole-pot mass for two representative units (left y-axis; upper curves) and the corresponding fertilizer-stock-tank solution depth (right y-axis; lower curves) for the 160 and 800 mg $L^{-1}$ nitrogen treatments. The series captures five consecutive nocturnal irrigation events. Vertical dashed red lines delimit the trigger window of an individual irrigation pulse; a rapid increase in lysimeter mass corresponds to a volumetric decrease in the respective treatment-supply tank. (b) Continuous monitoring of solution electrical conductivity (EC; left y-axis, dS $m^{-1}$) and pH (right y-axis) in the solution tanks during the same operational period. The stable baselines indicate consistent concentration control by the automated multichannel injection loops.

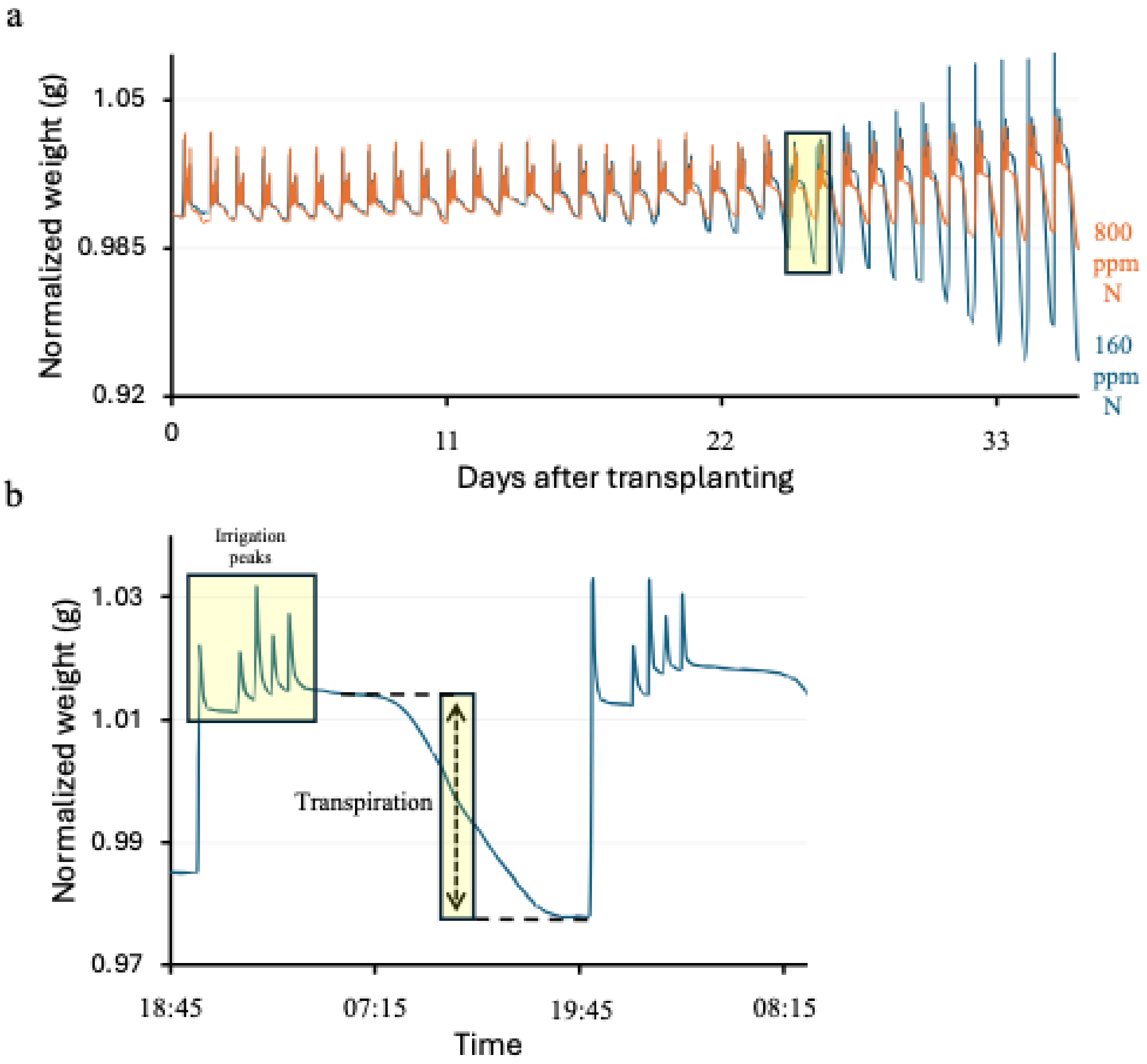


**Fig. S3.** High-resolution continuous gravimetric tracking of whole-pot mass dynamics and transpiration flux. Continuous physiological telemetry was recorded using an automated lysimeter array to monitor the real-time soil–plant–atmosphere water balance. (a) Longitudinal profiles of normalized pot mass (g) during cultivation for representative plants receiving the 160 and 800 mg $L^{-1}$ N treatments in Experiment 1 (Table 1 in the main manuscript). The increasing amplitude of the diurnal oscillations over time reflects canopy development and increasing daily transpiration. (b) High-resolution diurnal snapshot illustrating the noninvasive extraction of functional physiological traits from the gravimetric time series. The profile highlights (1) nocturnal irrigation peaks comprising five automated water-delivery pulses that restore the substrate to field capacity and (2) the steady daytime drawdown associated with hourly transpiration.

**Table S1.** Elemental composition and concentration gradients of the custom-formulated nutrient solutions used in the functional-phenotyping experiments. Fertigation treatments were designed to characterize physiological and ionomic responses across seven target-mineral gradients: nitrogen, potassium, phosphorus, calcium, magnesium, iron and molybdenum. Each experimental block comprised eight concentration levels (Treatment IDs 1–8), expressed in mg L$^{-1}$. Bold values indicate the target mineral varied within each experimental run. Variations in nontarget elements (e.g., sodium and sulfur) reflect counter-ion adjustments required to maintain electroneutrality and solution stability across the salt formulations.

| | | Mineral concentration (mg L$^{-1}$) | | | | | | | | | | | | |
|---|---|---|---|---|---|---|---|---|---|---|---|---|---|---|
| Target mineral | Treatment ID | N | K | P | Ca | Mg | Fe | Mn | Cu | Zn | Mo | Na | B | S |
| Nitrogen experiment | 1 | **20.0** | 80.3 | 30.0 | 80.5 | 30.3 | 2.7 | 1.8 | 0.1 | 0.3 | 0.1 | 1.1 | 0.03 | 68.7 |
| | 2 | **40.3** | 80.3 | 30.0 | 80.5 | 30.3 | 2.7 | 1.8 | 0.1 | 0.3 | 0.1 | 1.1 | 0.03 | 68.7 |
| | 3 | **80.0** | 80.3 | 30.0 | 80.5 | 30.3 | 2.7 | 1.8 | 0.1 | 0.3 | 0.1 | 1.1 | 0.03 | 68.7 |
| | 4 | **120.0** | 80.3 | 30.0 | 80.5 | 30.3 | 2.7 | 1.8 | 0.1 | 0.3 | 0.1 | 68.6 | 0.03 | 68.7 |
| | 5 | **160.3** | 80.3 | 30.0 | 80.5 | 30.3 | 2.7 | 1.8 | 0.1 | 0.3 | 0.1 | 98.3 | 0.03 | 68.7 |
| | 6 | **200.1** | 80.3 | 30.0 | 80.5 | 30.3 | 2.7 | 1.8 | 0.1 | 0.3 | 0.1 | 98.3 | 0.03 | 68.7 |
| | 7 | **400.1** | 80.3 | 30.0 | 80.5 | 30.3 | 2.7 | 1.8 | 0.1 | 0.3 | 0.1 | 98.3 | 0.03 | 68.7 |
| | 8 | **800.1** | 80.3 | 30.0 | 80.5 | 30.3 | 2.7 | 1.8 | 0.1 | 0.3 | 0.1 | 98.3 | 0.03 | 68.7 |
| Potassium experiment | 1 | 120.7 | **20.1** | 30.0 | 80.5 | 30.3 | 2.7 | 1.8 | 0.1 | 0.3 | 0.1 | 72.7 | 0.03 | 81.5 |
| | 2 | 120.3 | **50.2** | 30.0 | 80.5 | 30.3 | 2.7 | 1.8 | 0.1 | 0.3 | 0.1 | 53.8 | 0.03 | 81.5 |
| | 3 | 120.7 | **80.3** | 30.0 | 80.5 | 30.3 | 2.7 | 1.8 | 0.1 | 0.3 | 0.1 | 36.2 | 0.03 | 81.5 |
| | 4 | 120.6 | **100.4** | 30.0 | 80.5 | 30.3 | 2.7 | 1.8 | 0.1 | 0.3 | 0.1 | 24.1 | 0.03 | 81.5 |
| | 5 | 120.2 | **120.2** | 30.0 | 80.5 | 30.3 | 2.7 | 1.8 | 0.1 | 0.3 | 0.1 | 18.7 | 0.03 | 86.5 |
| | 6 | 120.2 | **160.5** | 30.0 | 80.5 | 30.3 | 2.7 | 1.8 | 0.1 | 0.3 | 0.1 | 18.7 | 0.03 | 103.0 |
| | 7 | 120.2 | **250.1** | 30.0 | 80.5 | 30.3 | 2.7 | 1.8 | 0.1 | 0.3 | 0.1 | 18.7 | 0.03 | 139.8 |
| | 8 | 120.2 | **500.1** | 30.0 | 80.5 | 30.3 | 2.7 | 1.8 | 0.1 | 0.3 | 0.1 | 18.7 | 0.03 | 242.5 |
| Phosphorus experiment | 1 | 120.2 | 100.5 | **5.1** | 80.8 | 30.3 | 2.7 | 1.8 | 0.1 | 0.3 | 0.1 | 8.1 | 0.03 | 77.0 |
| | 2 | 120.2 | 100.5 | **10.1** | 80.8 | 30.3 | 2.7 | 1.8 | 0.1 | 0.3 | 0.1 | 8.1 | 0.03 | 77.0 |
| | 3 | 120.2 | 100.5 | **20.1** | 80.8 | 30.3 | 2.7 | 1.8 | 0.1 | 0.3 | 0.1 | 8.1 | 0.03 | 77.0 |
| | 4 | 120.2 | 100.5 | **30.0** | 80.8 | 30.3 | 2.7 | 1.8 | 0.1 | 0.3 | 0.1 | 12.3 | 0.03 | 77.0 |
| | 5 | 120.2 | 100.1 | **40.0** | 80.8 | 30.3 | 2.7 | 1.8 | 0.1 | 0.3 | 0.1 | 12.3 | 0.03 | 71.7 |
| | 6 | 120.2 | 100.4 | **70.1** | 80.8 | 30.3 | 2.7 | 1.8 | 0.1 | 0.3 | 0.1 | 12.3 | 0.03 | 56.2 |
| | 7 | 120.4 | 100.8 | **100.2** | 80.8 | 30.3 | 2.7 | 1.8 | 0.1 | 0.3 | 0.1 | 30.4 | 0.03 | 59.5 |
| | 8 | 120.4 | 100.5 | **200.1** | 80.8 | 30.3 | 2.7 | 1.8 | 0.1 | 0.3 | 0.1 | 95.3 | 0.03 | 42.1 |
| Calcium experiment | 1 | 120.1 | 80.2 | 30.0 | **10.2** | 30.0 | 2.7 | 1.8 | 0.1 | 0.3 | 0.1 | 113.9 | 0.03 | 6.0 |
| | 2 | 120.1 | 80.0 | 30.0 | **25.4** | 30.0 | 2.7 | 1.8 | 0.1 | 0.3 | 0.1 | 113.9 | 0.03 | 18.2 |
| | 3 | 120.2 | 80.0 | 30.0 | **50.8** | 30.0 | 2.7 | 1.8 | 0.1 | 0.3 | 0.1 | 84.2 | 0.03 | 18.2 |
| | 4 | 120.1 | 80.0 | 30.0 | **80.4** | 30.0 | 2.7 | 1.8 | 0.1 | 0.3 | 0.1 | 49.1 | 0.03 | 18.2 |
| | 5 | 120.1 | 80.0 | 30.0 | **100.7** | 30.0 | 2.7 | 1.8 | 0.1 | 0.3 | 0.1 | 25.4 | 0.03 | 18.2 |
| | 6 | 120.1 | 80.0 | 30.0 | **140.5** | 30.2 | 2.7 | 1.8 | 0.1 | 0.3 | 0.1 | 0.0 | 0.03 | 32.8 |
| | 7 | 119.2 | 80.0 | 30.0 | **250.4** | 30.1 | 2.7 | 1.8 | 0.1 | 0.3 | 0.1 | 0.0 | 0.03 | 120.1 |
| | 8 | 119.2 | 80.0 | 30.0 | **375.0** | 30.1 | 2.7 | 1.8 | 0.1 | 0.3 | 0.1 | 0.0 | 0.03 | 206.6 |
| Magnesium experiment | 1 | 120.7 | 80.3 | 30.0 | 80.5 | **5.1** | 2.7 | 1.8 | 0.1 | 0.3 | 0.1 | 45.7 | 0.03 | 48.3 |
| | 2 | 120.7 | 80.3 | 30.0 | 80.5 | **10.1** | 2.7 | 1.8 | 0.1 | 0.3 | 0.1 | 45.7 | 0.03 | 54.9 |
| | 3 | 120.7 | 80.3 | 30.0 | 80.5 | **20.2** | 2.7 | 1.8 | 0.1 | 0.3 | 0.1 | 45.7 | 0.03 | 68.2 |
| | 4 | 120.7 | 80.3 | 30.0 | 80.5 | **30.3** | 2.7 | 1.8 | 0.1 | 0.3 | 0.1 | 45.7 | 0.03 | 81.5 |
| | 5 | 120.2 | 80.3 | 30.0 | 80.5 | **50.0** | 2.7 | 1.8 | 0.1 | 0.3 | 0.1 | 10.6 | 0.03 | 81.5 |
| | 6 | 120.2 | 80.4 | 30.0 | 80.5 | **100.3** | 2.7 | 1.8 | 0.1 | 0.3 | 0.1 | 45.7 | 0.03 | 94.6 |
| | 7 | 120.7 | 80.3 | 30.0 | 80.5 | **200.9** | 2.7 | 1.8 | 0.1 | 0.3 | 0.1 | 10.6 | 0.03 | 202.5 |
| | 8 | 120.7 | 80.3 | 30.0 | 80.5 | **400.9** | 2.7 | 1.8 | 0.1 | 0.3 | 0.1 | 10.6 | 0.03 | 465.9 |
| Iron experiment | 1 | 120.1 | 80.3 | 30.0 | 80.5 | 30.3 | **0.5** | 1.8 | 0.1 | 0.3 | 0.1 | 45.9 | 0.03 | 81.5 |
| | 2 | 120.1 | 80.3 | 30.0 | 80.5 | 30.3 | **0.9** | 1.8 | 0.1 | 0.3 | 0.1 | 45.9 | 0.03 | 81.5 |
| | 3 | 120.1 | 80.3 | 30.0 | 80.5 | 30.3 | **1.8** | 1.8 | 0.1 | 0.3 | 0.1 | 45.9 | 0.03 | 81.5 |
| | 4 | 120.1 | 80.3 | 30.0 | 80.5 | 30.3 | **2.7** | 1.8 | 0.1 | 0.3 | 0.1 | 45.9 | 0.03 | 81.5 |
| | 5 | 120.1 | 80.3 | 30.0 | 80.5 | 30.3 | **3.6** | 1.8 | 0.1 | 0.3 | 0.1 | 45.9 | 0.03 | 81.5 |
| | 6 | 120.1 | 80.3 | 30.0 | 80.5 | 30.3 | **4.6** | 1.8 | 0.1 | 0.3 | 0.1 | 45.9 | 0.03 | 81.5 |
| | 7 | 120.1 | 80.3 | 30.0 | 80.5 | 30.3 | **9.1** | 1.8 | 0.1 | 0.3 | 0.1 | 45.9 | 0.03 | 81.5 |
| | 8 | 120.1 | 80.3 | 30.0 | 80.5 | 30.3 | **18.2** | 1.8 | 0.1 | 0.3 | 0.1 | 45.9 | 0.03 | 81.5 |
| Molybdenum experiment | 1 | 120.8 | 80.3 | 30.0 | 80.4 | 30.3 | 2.7 | 1.8 | 0.1 | 0.3 | **0.01** | 40.5 | 0.03 | 72.6 |
| | 2 | 120.8 | 80.3 | 30.0 | 80.4 | 30.3 | 2.7 | 1.8 | 0.1 | 0.3 | **0.04** | 40.5 | 0.03 | 72.6 |
| | 3 | 120.8 | 80.3 | 30.0 | 80.4 | 30.3 | 2.7 | 1.8 | 0.1 | 0.3 | **0.08** | 40.5 | 0.03 | 72.6 |
| | 4 | 120.8 | 80.3 | 30.0 | 80.4 | 30.3 | 2.7 | 1.8 | 0.1 | 0.3 | **0.12** | 40.5 | 0.03 | 72.6 |
| | 5 | 120.8 | 80.3 | 30.0 | 80.4 | 30.3 | 2.7 | 1.8 | 0.1 | 0.3 | **0.14** | 40.5 | 0.03 | 72.6 |
| | 6 | 120.8 | 80.3 | 30.0 | 80.4 | 30.3 | 2.7 | 1.8 | 0.1 | 0.3 | **1** | 40.5 | 0.03 | 72.6 |
| | 7 | 120.8 | 80.3 | 30.0 | 80.4 | 30.3 | 2.7 | 1.8 | 0.1 | 0.3 | **5** | 40.5 | 0.03 | 72.6 |
| | 8 | 120.8 | 80.3 | 30.0 | 80.4 | 30.3 | 2.7 | 1.8 | 0.1 | 0.3 | **10** | 40.5 | 0.03 | 72.6 |

**Table. S2** Global modeling architecture, training hyperparameters, and optimized target-specific configurations for the xLSTM microclimatic imputation networks. Deep learning recurrent neural networks were developed within the PyTorch framework, utilizing the Extended Long Short-Term Memory (xLSTM) library, to reconstruct high-resolution 3-minute environmental data streams for the compromised Potassium Chinese broccoli cultivation cycle. The computational foundation for all pipelines utilized a uniform structural sequence: a linear input projection feeding a stack of matrix-memory Extended Long Short-Term Memory (mLSTM) blocks, paired with an intermediate dropout regularization layer and a single-node dense linear output layer, executing one specialized model independently per target variable. Prior to model training, raw environmental time-series data were passed through a prespecified, physically-motivated physical-range quality-control filter (VPD ≤ 6 kPa and Ambient temperature ≤ 45 C) to exclude sensor anomalies and subsequently normalized using MinMaxScaler applied independently per feature and target; each scaler was fitted on the training partition only and then applied to the validation and test partitions to prevent normalization leakage. Chronological partitioning was enforced across the time series array to prevent temporal data leakage, utilizing a strict 70%/15%/15% split for training, validation, and testing respectively. All network optimization loops were driven by the AdamW optimizer (learning rate $1x10^{-3}$, weight decay $1x10^{-4}$) minimizing a Mean squared error (MSE) loss function. Structural search boundaries were initially mapped via an exhaustive grid search and subsequently refined through per-target Bayesian optimization (Optuna Tree-structured Parzen Estimator, up to 60 independent tuning trials each with median pruning) alongside a sequence-length (lookback) sweep, and final weights were optimized with an early-stopping patience threshold of 6 epochs. To ensure strict experimental reproducibility, each configuration was retrained across five independent random seeds (42, 1, 7, 21, 99). Optimized target specific hyperparameters alongside out-of-sample test performance metrics (Coefficient of Determination, $R^2$; root mean squared error, RMSE; and mean absolute error, MAE), reported as mean ± standard deviation across seeds, are structured below. Predicted-versus-observed and diurnal trajectory figures depict, for each target, the representative run whose test $R^2$ is closest to the corresponding five-seed mean, annotated with these same mean ± standard deviation values.

| Target environmental feature | Lookback (steps) | Embedded dimension | mLSTM blocks | Dropout rate | Test $R^2$ | Test RMSE | Test MAE |
|---|---|---|---|---|---|---|---|
| PPFD (μmol $m^{-2}$ $s^{-1}$) | 30 | 64 | 3 | 0.0 | 0.621±0.003 | 130.93±0.51 | 62.87±2.38 |
| RH (%) | 5 | 64 | 3 | 0.2 | 0.888±0.008 | 5.33±0.18 | 3.81±0.2 |
| Temp (°C) | 30 | 32 | 4 | 0.2 | 0.870±0.007 | 1.62±0.04 | 1.04±0.03 |
| VPD (kPa) | 5 | 96 | 3 | 0.2 | 0.813±0.002 | 0.46±0.01 | 0.28±0.01 |

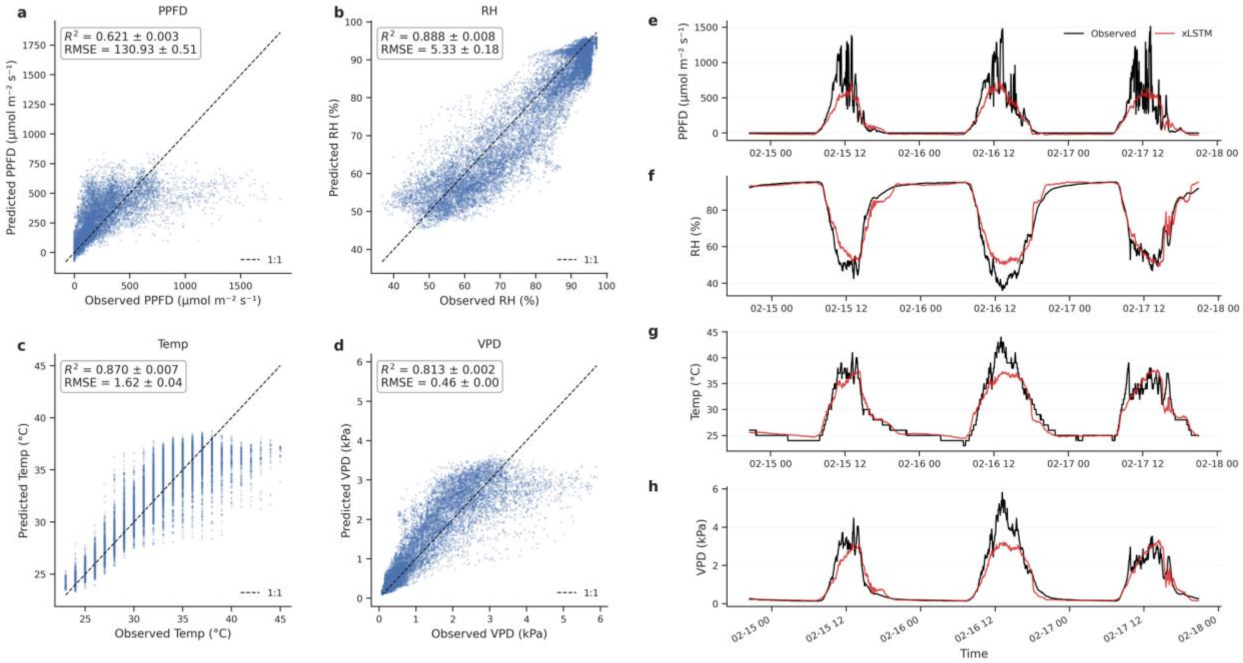


**Fig. S4** Out-of-sample predictive performance and time-series alignment of the deep learning xLSTM microclimatic reconstruction pipeline. Recurrent neural networks were trained to model the non-linear thermodynamic relationships between external ambient weather data and the greenhouse microclimate interior. (a-d) scatter plots illustrating the out-of-sample predictive accuracy against true internal sensor observation across the held-out test dataset for: (a) Photosynthetic photon flux density (PPFD; µmol $m^{-2}$ $s^{-1}$), (b) Relative humidity (RH; %), (c) Ambient temperature (Temp; C), and (d) Vapor pressure deficit (VPD; kPa). Inset text boxes report the coefficient of determination ($R^2$) and root mean square error (RMSE) for each environmental target, expressed as the mean ± standard deviation across the five independently seeded training runs; panels depict the representative run whose test R2 is closest to the corresponding mean. The black dashed line indicates the 1:1 line of perfect alignment. (e-h) Continuous 72-hour test-set trajectories contrasting observed sensor recordings (black line) against the xLSTM-reconstructed predictions (red line) for (e) PPFD, (f) RH, (g) Temp, and (h) VPD. The close alignment in phase and amplitude across diurnal photoperiod peaks and nocturnal baselines - most pronounced for relative humidity, temperature, and vapour pressure deficit, with PPFD faithfully tracking the daytime photoperiod envelope - demonstrates the structural robustness of the imputation pipeline used to recover missing feature matrices for the compromised Chinese broccoli experiment.

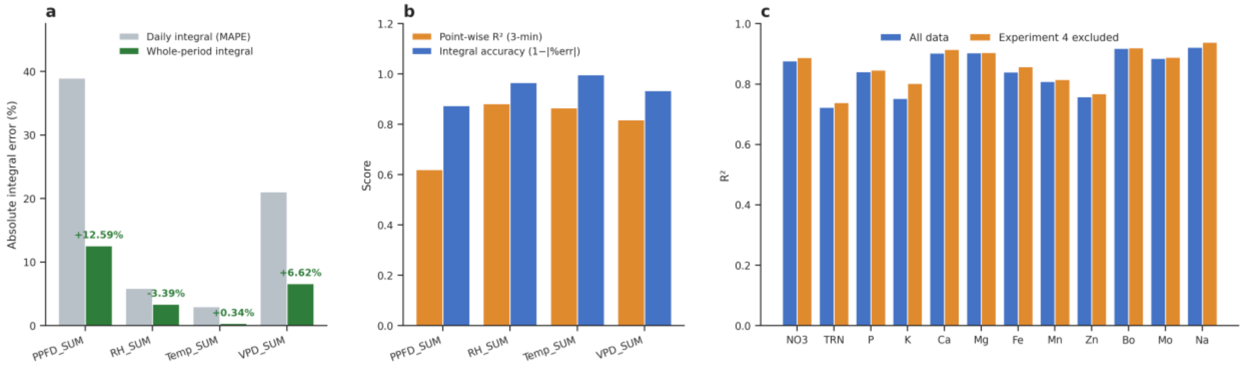


**Fig. S5** Accuracy of the xLSTM microclimate imputation for Experiment 4 and its effect on ionomic prediction. (a) Absolute error of the cumulative environmental integrals reconstructed by the xLSTM, evaluated on the final-15% chronological hold-out spanning 12 February to 19 June 2025 (~46,700 three-minute records retained after quality filtering, not seen during training). Cumulative features were computed as trapezoidal time-integrals of the 3-min series over the diurnal photoperiod (07:00-19:00), consistent with the feature definition in Section 2.3.2 (PPFD_SUM, mol $m^{-2}$; RH_SUM, % d; Temp_SUM, C d; VPD_SUM, kPa d); the reported errors are percentage errors and are therefore invariant to the integration constant and unit choice. Grey bars, mean absolute percentage error of the per-day integrals (MAPE); green bars, signed percentage error of the integral accumulated over the whole hold-out. Values above the green bars give the whole-window error (PPFD_SUM +12.6%, RH_SUM -3.4%, Temp_SUM +0.3%, VPD_SUM +6.6%; signed). Panels a and b depict the representative run (the seed shown throughout Fig. S4). (b) Point-wise reconstruction skill (orange; R2 of the 3-min predicted vs measured series) versus cumulative-integral accuracy (blue; 1 - |whole-window % error|) for the same four features. (c) Leakage-free cross-validated performance (TabPFN v3, pooled out-of-fold R2 under stratified round-robin DOB-SCV cross-validation; Fig. S6) for all twelve harvested-leaf minerals, trained on all data (blue) versus with the xLSTM-imputed Experiment 4 removed (orange). Mean $|dR^2|$ = 0.013 across targets; mean $R^2$ 0.844 to 0.857; the largest change is an increase for potassium (K, the mineral gradient of Experiment 4).

**Supplementary Note. 1** Analysis of Radiative vs Thermodynamic Predictive Discrepancies under Tropical Atmospheric Conditions. The structural variations in out-of-sample accuracy across the four microclimatic targets reflect the distinct spatial-temporal scales governing equatorial atmospheric phenomena. While regional ambient temperature ($R^2$=0.870), relative humidity ($R^2$=0.888), and VPD ($R^2$=0.813) display high thermodynamic uniformity and strong macro-scale coupling over 5-km length, the PPFD ($R^2$=0.621) is dictated by highly localized, stochastic environmental boundary conditions.

In tropical equatorial marine-influenced climates, intermittent convective cloud formations produce rapid, high-frequency, micro-scale fluctuations in solar radiance. Because these transient cloud transitions occur directly over the primary greenhouse canopy matrix, they often do not align chronologically or spatially with the sensor records logged at the external meteorological station 5 km away. This spatial decoupling imposes a structural mathematical ceiling on point-to-point time-series correlations ($R^2$) and inflates the instantaneous root mean square error (RMSE=130.93 µmol $m^{-2}$ $s^{-1}$) for radiative metrics, even under highly optimized network weight configurations.

Crucially, however, the tight phase synchronization and preserved diurnal envelope captured across the longitudinal trajectories (Fig. S4e) confirm that the xLSTM network successfully reconstructed the essential day-length envelopes and integrated photoperiod boundaries. This ensures that when these high-resolution data arrays are integrated over the life cycle duration to engineer the primary cumulative macro-features for the downstream pipelines, the thermodynamic cumulative drivers (Temp_SUM, RH_SUM) remain highly accurate, while the cumulative light driver (PPFD_SUM) preserves the correct photoperiod envelope with a bounded, quantifiable magnitude bias (quantified below).

To move this argument from qualitative to quantitative, we evaluated the imputation on the model's chronological hold-out spanning 12 February to 19 June 2025 (~46,700 three-minute records retained after quality filtering and withheld from training), integrating the predicted and measured internal series over time to reconstruct the cumulative features actually consumed downstream. The point-wise reconstruction skill ($R^2$ = 0.621, 0.888, 0.870 and 0.813 for PPFD, relative humidity, temperature and VPD) propagated to the integrated drivers in a strongly target-dependent manner. Integrating over the diurnal photoperiod (07:00-19:00) as in the main-text feature definition, the thermodynamically well-constrained variables largely cancelled their high-frequency, sign-random point-wise deviations under temporal integration, and the cumulative integrals were recovered with small whole-window errors of -3.4 % (RH_SUM) and +0.3 % (Temp_SUM). For the radiative and radiation-coupled variables the point-wise deviations were more systematically signed and therefore accumulated rather than cancelled, leaving residual whole-window biases of +12.6 % (PPFD_SUM) and +6.6 % (VPD_SUM) (Fig. S5a,b). The residual PPFD_SUM bias is the integral signature of the structurally bounded instantaneous PPFD correlation described above: where the external station cannot resolve the micro-scale convective cloud transitions occurring over the canopy, the resulting point-wise errors retain a net sign and propagate into the integrated light driver rather than averaging out. The engineered thermodynamic macro-features are therefore physically faithful to the true cultivation history, whereas the cumulative light driver carries a quantified positive bias that should be acknowledged when interpreting PPFD-dependent effects.

Finally, to verify that the single imputed experiment does not itself drive the reported predictive performance, we repeated the leakage-free cross-validation (stratified round-robin DOB-SCV cross-validation, Fig. S6; TabPFN v3) with Experiment 4 (the potassium gradient) removed. Per-target out-of-fold $R^2$ changed by only 0.013 on average (mean 0.844 to 0.857), with every target remaining within $|\Delta R^2| \leq 0.05$ and the largest single change being a slight increase for potassium - the mineral gradient defined by that experiment (Fig. S5c). Because Experiment 4's internal microclimate is by construction unmeasured, the integral-error analysis is performed on the hold-out as a proxy that reproduces the identical external-to-internal reconstruction task; together the two analyses show that the imputation is accurate at the integral level and that the study's conclusions are insensitive to the inclusion of the imputed experiment.

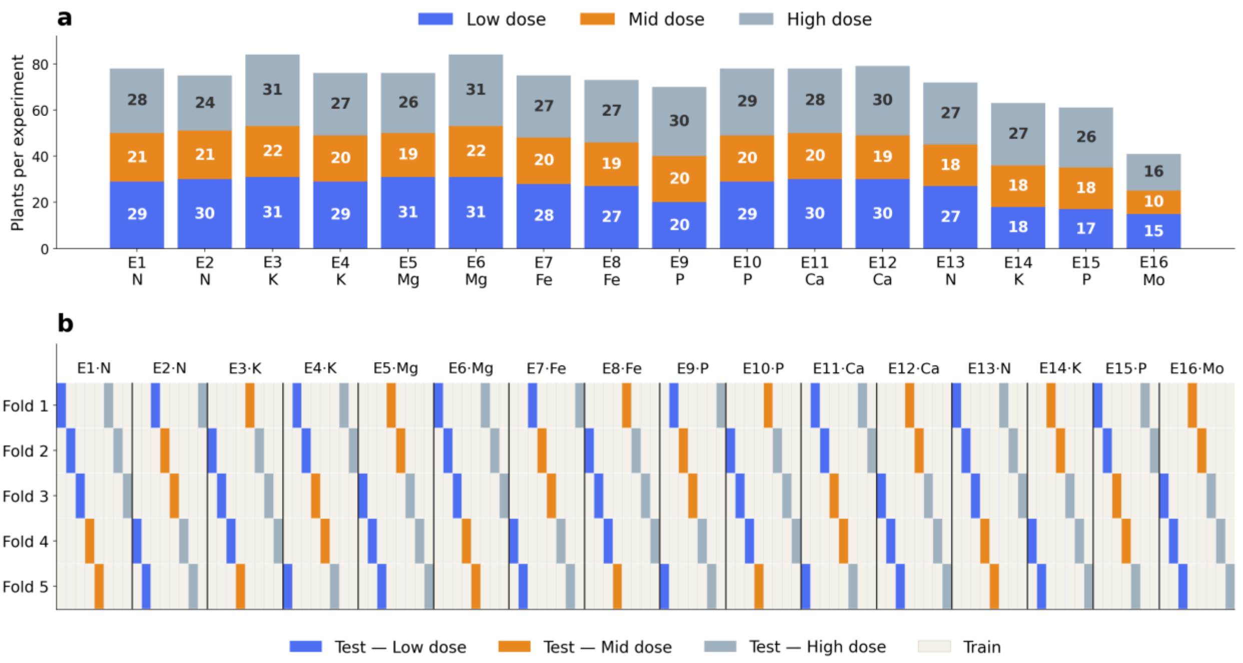


**Fig. S6** Schematic architecture of the stratified round-robin (DOB-SCV) cross-validation strategy used for all mineral-prediction model evaluation in this study. The 16 independent cultivation experiments were first decomposed into complete dosing conditions, where each condition comprises all biological replicates sharing the same experiment, target-mineral gradient and concentration tier. Each dosing condition was treated as a single non-divisible group, preventing plants grown under the same nutrient-dose condition from being split between the training and validation sets. Folds were built by a single deterministic distribution-balancing procedure (Distribution-Optimally-Balanced Stratified Cross-Validation, DOB-SCV): within each dose tier, complete dosing-condition groups were distributed across the five folds in round-robin order, so that every fold receives a representative, dose-balanced slice of the full design and every experiment is represented in every fold. Panel a shows the resulting per-experiment dose composition; Panel (b) shows the fold assignment, where blue, orange and grey blocks mark conditions held out for testing at the Low, Mid and High dose tiers respectively, and beige blocks denote conditions used for training. This construction yields complete experiment coverage by design, comparable dose-tier composition across folds (condition-level Pearson's $\chi^2$ test, $p>0.999$; 24-26 dosing conditions per validation fold) and no same-condition leakage (group-disjoint folds; permuted-label R-squared approximately 0), providing the single deterministic partition on which all cross-validated mineral-prediction metrics reported in this study are computed.

**Table S3.** AutoGluon configuration and ensemble structure used for automated machine-learning benchmarking.

| Parameter | Value |
|---|---|
| Software | AutoGluon |
| Version | 1.5.0 |
| Module | autogluon.tabular |
| Predictor | TabularPredictor |
| Problem type | Regression |
| Evaluation metric | Root mean squared error |
| Preset | best_quality |
| Bagging folds | 8 |
| Bagging repeats | 1 |
| Maximum stacking level | 3 |
| Training budget | 3600 seconds per fit; 900 seconds at reduced sizes (learning curves) |
| Random seed | 42 (training-subsample draw); AutoGluon internals unseeded |
| Base learners | LightGBM, LightGBM-XT, LightGBM-Large, XGBoost, CatBoost, Random Forest, Extra Trees, NeuralNetTorch, NeuralNetFastAI |
| Final ensemble | WeightedEnsemble |
| Ensemble selection | Greedy weighted ensemble |

**Table S4.** AutoGluon 1.5 stacked-ensemble architecture for each leaf-mineral target. All targets used the best_quality preset: an eightfold-bagged, multilayer stacked ensemble optimized for RMSE and fitted independently within each of the five Distribution-Optimally-Balanced Stratified Cross-Validation (DOB-SCV) folds (Section 2.5.2 of the main manuscript). "Max stacking level" is the deepest layer reached in any fold (level 1, base learners fitted to the original features; levels 2–3, stacker layers and the weighted-ensemble head). "Base models" is the mean number of candidate learners trained per fold. "Ensemble members" is the mean number of learners assigned nonzero weight in the final greedy weighted ensemble. Both values were averaged across the five folds. "Top families" identifies the two base-learner families with the greatest summed ensemble weights.

| Target | Bagging folds | Max stacking level | Base models (mean) | Ensemble members (mean) | Top base-learner families |
|---|---|---|---|---|---|
| $NO_3^-$ | 8 | 3 | 94 | 4 | NeuralNetTorch, NeuralNetFastAI |
| TRN | 8 | 3 | 78 | 7 | LightGBM, NeuralNetTorch |
| P | 8 | 3 | 88 | 6 | NeuralNetTorch, ExtraTrees |
| K | 8 | 3 | 90 | 6 | NeuralNetTorch, NeuralNetFastAI |
| Ca | 8 | 3 | 97 | 7 | NeuralNetTorch, ExtraTreesMSE |
| Mg | 8 | 3 | 87 | 7 | ExtraTrees, NeuralNetTorch |
| Fe | 8 | 3 | 121 | 4 | NeuralNetTorch, LightGBM |
| Mn | 8 | 3 | 104 | 7 | LightGBM, NeuralNetTorch |
| Zn | 8 | 3 | 95 | 6 | NeuralNetTorch, NeuralNetFastAI |
| B | 8 | 3 | 91 | 4 | NeuralNetTorch, NeuralNetFastAI |
| Mo | 8 | 3 | 63 | 5 | NeuralNetTorch, CatBoost |
| Na | 8 | 3 | 102 | 6 | NeuralNetTorch, NeuralNetFastAI |

**Table S5.** TabPFN regression configuration used for tabular foundation-model benchmarking. The table summarizes the implementation settings used to evaluate TabPFN v3 as a regressor for plant-ionomic prediction, including the software version, model checkpoint, inference-ensemble size, preprocessing mode, precision settings, hardware backend (NVIDIA L40S GPUs on the HURCS cluster), parallelization and random seed.

| Parameter | Value |
|---|---|
| Python package | tabpfn v8.0.3 |
| Model family | TabPFN v3 regressor |
| Checkpoint | tabpfn-v3-regressor-v3_default |
| n_estimators | 32 |
| softmax_temperature | 0.9 (default) |
| average_before_softmax | False (default) |
| inference_precision | Automatic (default) |
| fit_mode | fit_preprocessors (default) |
| ignore_pretraining_limits | True |
| device | CUDA |
| n_jobs | 8 |
| random_state | 42 |

**Table S6.** Mean Spearman rank correlation (ρ) across four representative targets (total reduced nitrogen, Ca, Fe and Mo) between per-feature mean absolute SHAP vectors obtained under each pair of settings. All other settings were held at the robustness-analysis baseline: 150 explained instances, 2,048 coalitions, k = 5 and the first Distribution-Optimally-Balanced Stratified Cross-Validation (DOB-SCV) fold. Lower ρ for cross-validation-fold comparisons reflects the grouped design, in which each fold withholds a disjoint set of dosing conditions. A value of ρ = 1 indicates an unchanged feature ranking.

| Configuration | Setting comparison | TabPFN v3 (ρ) | AutoGluon 1.5 (ρ) |
|---|---|---|---|
| Coalition count | 1024 vs 2048 | 0.995 | 0.998 |
| Background size (k) | 5 vs 10 | 0.953 | 0.983 |
| | 5 vs 20 | 0.955 | 0.969 |
| | 10 vs 20 | 0.981 | 0.983 |
| Explained-sample set | A vs B | 0.979 | 0.958 |
| Cross-validation fold | 1 vs 2 | 0.929 | 0.905 |
| | 0 vs 1 | 0.933 | 0.943 |
| | 0 vs 2 | 0.918 | 0.870 |
